# Type 2 and interferon inflammation strongly regulate SARS-CoV-2 related gene expression in the airway epithelium

**DOI:** 10.1101/2020.04.09.034454

**Authors:** Satria P. Sajuthi, Peter DeFord, Nathan D. Jackson, Michael T. Montgomery, Jamie L. Everman, Cydney L. Rios, Elmar Pruesse, James D. Nolin, Elizabeth G. Plender, Michael E. Wechsler, Angel CY Mak, Celeste Eng, Sandra Salazar, Vivian Medina, Eric M. Wohlford, Scott Huntsman, Deborah A. Nickerson, Soren Germer, Michael C. Zody, Gonçalo Abecasis, Hyun Min Kang, Kenneth M. Rice, Rajesh Kumar, Sam Oh, Jose Rodriguez-Santana, Esteban G. Burchard, Max A. Seibold

## Abstract

Coronavirus disease 2019 (COVID-19) outcomes vary from asymptomatic infection to death. This disparity may reflect different airway levels of the SARS-CoV-2 receptor, ACE2, and the spike protein activator, TMPRSS2. Here we explore the role of genetics and co-expression networks in regulating these genes in the airway, through the analysis of nasal airway transcriptome data from 695 children. We identify expression quantitative trait loci (eQTL) for both *ACE2* and *TMPRSS2*, that vary in frequency across world populations. Importantly, we find *TMPRSS2* is part of a mucus secretory network, highly upregulated by T2 inflammation through the action of interleukin-13, and that interferon response to respiratory viruses highly upregulates *ACE2* expression. Finally, we define airway responses to coronavirus infections in children, finding that these infections upregulate *IL6* while also stimulating a more pronounced cytotoxic immune response relative to other respiratory viruses. Our results reveal mechanisms likely influencing SARS-CoV-2 infectivity and COVID-19 clinical outcomes.

## Introduction

In December of 2019, a novel Coronavirus, SARS-CoV-2, emerged in China and has gone on to trigger a global pandemic of Coronavirus Disease 2019 (COVID-19), the respiratory illness caused by this virus^1^. While most individuals with COVID-19 experience mild cold symptoms (cough and fever), some develop more severe disease including pneumonia, which often necessitates mechanical ventilation^2^. In fact, an estimated 5.7% of COVID-19 illnesses are fatal^3^. Enhanced risk of poor outcomes for COVID-19 has been associated with a number of factors including advanced age, male sex, and underlying cardiovascular and respiratory conditions^4, 5^. Yet, while the majority of serious COVID-19 illness occurs in adults over 60, children are also thought to be highly susceptible to infection. Moreover, recent data suggest that 38% of COVID-19 cases occurring in children are of moderate severity and 5.8% are severe or critical^6^, highlighting a need for studying risk factors of illness in this population as well.

One factor that may underlie variation in clinical outcomes of COVID-19 is the extent of gene expression in the airway of the SARS-CoV-2 entry receptor, *ACE2*, and *TMPRSS2*, the host protease that cleaves the viral spike protein and thus allows for efficient virus-receptor binding^7^. Expression of these genes and their associated programs in the nasal airway epithelium is of particular interest given that the nasal epithelium is the primary site of infection for upper airway respiratory viruses, including coronaviruses, and acts as the gateway through which upper airway infections can spread into the lung. The airway epithelium is composed of multiple resident cell types (e.g., mucus secretory, ciliated, basal stem cells, and rare epithelial cell types) interdigitated with immune cells (e.g. T cells, mast cells, macrophages), and the relative abundance of these cell types in the epithelium can greatly influence the expression of particular genes^8-10^, including *ACE2* and *TMPRSS2*. Furthermore, since the airway epithelium acts as a sentinel for the entire respiratory system, its cellular composition, along with its transcriptional and functional characteristics, are significantly shaped by interaction with environmental stimuli. These stimuli may be inhaled (e.g., cigarette smoke, allergens, microorganisms) or endogenous, such as when signaling molecules are produced by airway immune cells present during different disease states. One such disease state is allergic airway inflammation caused by type 2 (T2) cytokines (IL-4, IL-5, IL-13), which is common in both children and adults and has been associated with the development of both asthma and COPD in a subgroup of patients^11-13^. T2 cytokines are known to greatly modify gene expression in the airway epithelium, both through transcriptional changes within cells and epithelial remodeling in the form of mucus metaplasia^11, 14, 15^. Microbial infection is another strong regulator of airway epithelial expression. In particular, respiratory viruses can modulate the expression of thousands of genes within epithelial cells, while also recruiting and activating an assortment of immune cells^16-18^. Even asymptomatic nasal carriage of respiratory viruses, which is especially common in childhood, has been shown to be associated with both genome-wide transcriptional re-programming and infiltration of macrophages and neutrophils in the airway epithelium^19^, demonstrating how viral infection can drive pathology even without overt signs of illness.

Genetic variation is another factor that may regulate gene expression in the airway epithelium. Indeed, expression quantitative trait loci (eQTL) analyses carried out in many tissues have suggested that as many as 70% of genes expressed by a tissue or organ are under genetic control^20^. Severity of human rhinovirus (HRV) respiratory illness has specifically been associated with genetic variation in the epithelial genes *CDHR3*^21^ and the *ORMDL3*^22^ and, given differences in genetic variation across world populations, it is possible that functional genetic variants in SARS-CoV-2-related genes could partly explain population differences in COVID-19 clinical outcomes.

Finally, there are important questions regarding the host response to SARS-CoV-2 infection. For example, it is unclear whether specific antiviral defenses in the epithelium are blocked by SARS-CoV-2 or whether the virus may trigger epithelial or immune cell pathways that prolong airway infection, and/or even incite a hyperinflammatory state in the lungs in some individuals that leads to more severe disease. Although large cohorts of subjects infected by the novel coronavirus are still lacking, much can be learned by exploring transcriptional responses to other coronavirus strains. In particular, because nasal airway brushings capture both epithelial and immune cells present at the airway surface, such samples collected from a cohort of subjects infected by a range of viruses provide an opportunity to comprehensively investigate the potentially varied and cascading effects of coronavirus infection on airway expression and function.

In this study, we first use single cell RNA-sequencing (scRNA-seq) to elucidate the cellular distribution of *ACE2* and *TMPRSS2* expression in the nasal airway epithelium. We also perform network and eQTL analysis of bulk gene expression data on nasal airway epithelial brushings collected from a large cohort of asthmatic and healthy children in order to identify the genetic and biological regulatory mechanisms governing *ACE2* and *TMPRSS2* expression. We then use multi-variable modeling to estimate the relative contribution of these factors to population variation in the expression of these two genes, and by performing experiments on mucociliary airway epithelial cultures confirm a dominant role for both T2 inflammation and viral infection in regulating expression of *ACE2* and *TMPRSS2*. Finally, we define the cellular and transcriptional responses to *in vivo* coronavirus infections in the nasal airway of children.

## Results

### *ACE2* and *TMPRSS2* are expressed by multiple nasal airway epithelial cell types

We first examined *ACE2* and *TMPRSS2* expression at a cell type level through single cell RNA sequencing (scRNA-seq) of a nasal airway epithelial brushing from an asthmatic subject. Shared Nearest Neighbor (SNN)-based clustering of 8,291 cells identified 9 epithelial and 3 immune cell populations (Figure 1a, Supplementary Table 1). We found that 7 epithelial cell populations contained *ACE2*^*+*^ cells (at low frequency), with the highest frequency of positive cells found among basal/early secretory cells, ciliated cells, and secretory cells (Figure 1b). We did not observe meaningful *ACE2* expression among any of the immune cell populations, which included T cells, dendritic cells, and mast cells. We found *TMPRSS2* to be expressed by all epithelial cell types, with a higher frequency of positive cells among the different cell types, compared to *ACE2* (Figure 1b,c). A small number of mast cells were also *TMPRSS2*^+^ (Figure 1c).

**Table 1.**
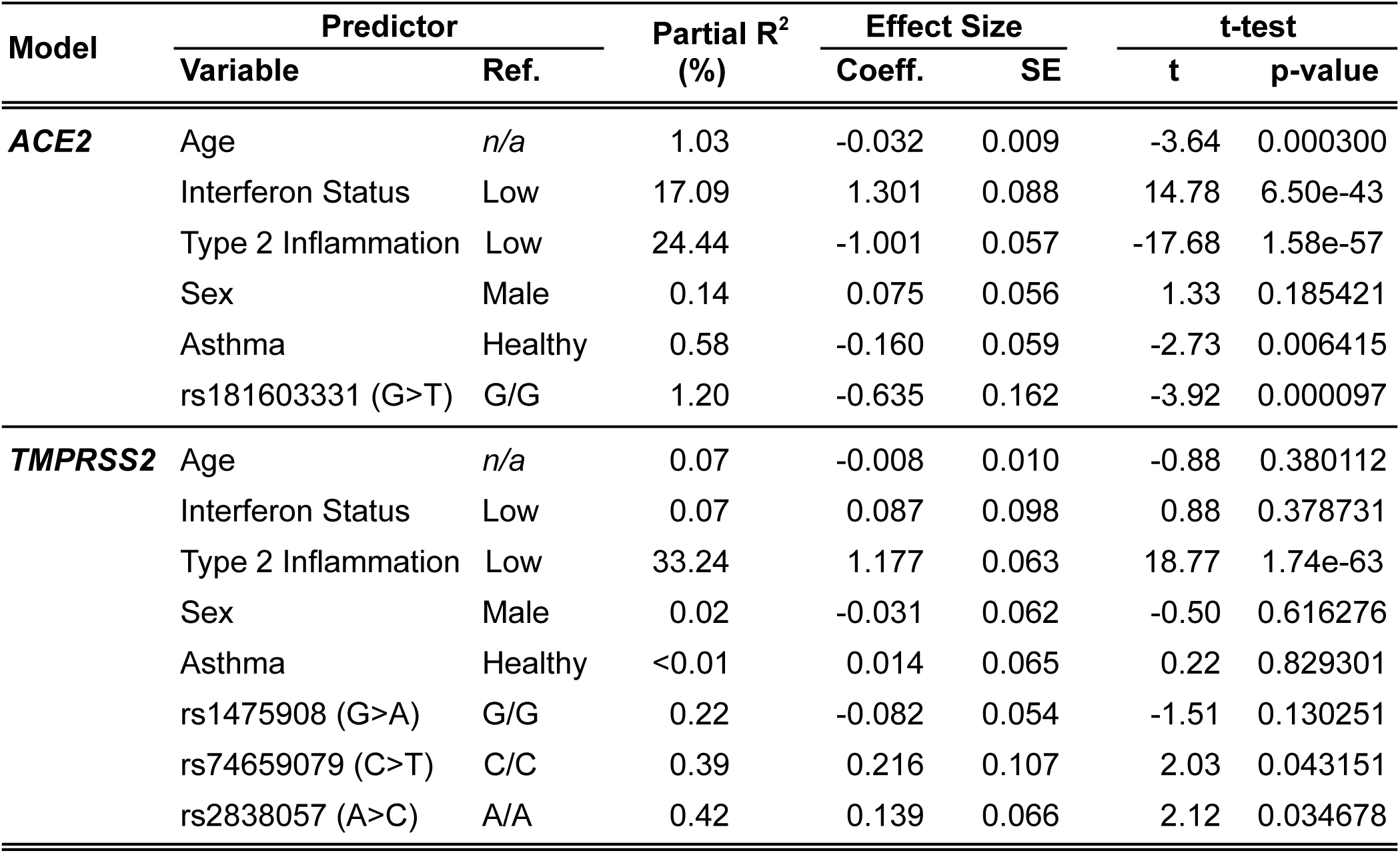
Results for multivariate models predicting ACE2 and TMPRSS2 expression.

| Model | Predictor |  | Partial R<sup>2</sup><br>(%) | Effect Size |  | t-test |  |
| --- | --- | --- | --- | --- | --- | --- | --- |
|  | Variable | Ref. |  | Coeff. | SE | t | p-value |
| **ACE2** | Age | *n/a* | 1.03 | -0.032 | 0.009 | -3.64 | 0.000300 |
|  | Interferon Status | Low | 17.09 | 1.301 | 0.088 | 14.78 | 6.50e-43 |
|  | Type 2 Inflammation | Low | 24.44 | -1.001 | 0.057 | -17.68 | 1.58e-57 |
|  | Sex | Male | 0.14 | 0.075 | 0.056 | 1.33 | 0.185421 |
|  | Asthma | Healthy | 0.58 | -0.160 | 0.059 | -2.73 | 0.006415 |
|  | rs181603331 (G>T) | G/G | 1.20 | -0.635 | 0.162 | -3.92 | 0.000097 |
| **TMPRSS2** | Age | *n/a* | 0.07 | -0.008 | 0.010 | -0.88 | 0.380112 |
|  | Interferon Status | Low | 0.07 | 0.087 | 0.098 | 0.88 | 0.378731 |
|  | Type 2 Inflammation | Low | 33.24 | 1.177 | 0.063 | 18.77 | 1.74e-63 |
|  | Sex | Male | 0.02 | -0.031 | 0.062 | -0.50 | 0.616276 |
|  | Asthma | Healthy | <0.01 | 0.014 | 0.065 | 0.22 | 0.829301 |
|  | rs1475908 (G>A) | G/G | 0.22 | -0.082 | 0.054 | -1.51 | 0.130251 |
|  | rs74659079 (C>T) | C/C | 0.39 | 0.216 | 0.107 | 2.03 | 0.043151 |
|  | rs2838057 (A>C) | A/A | 0.42 | 0.139 | 0.066 | 2.12 | 0.034678 |

**Figure 1.**
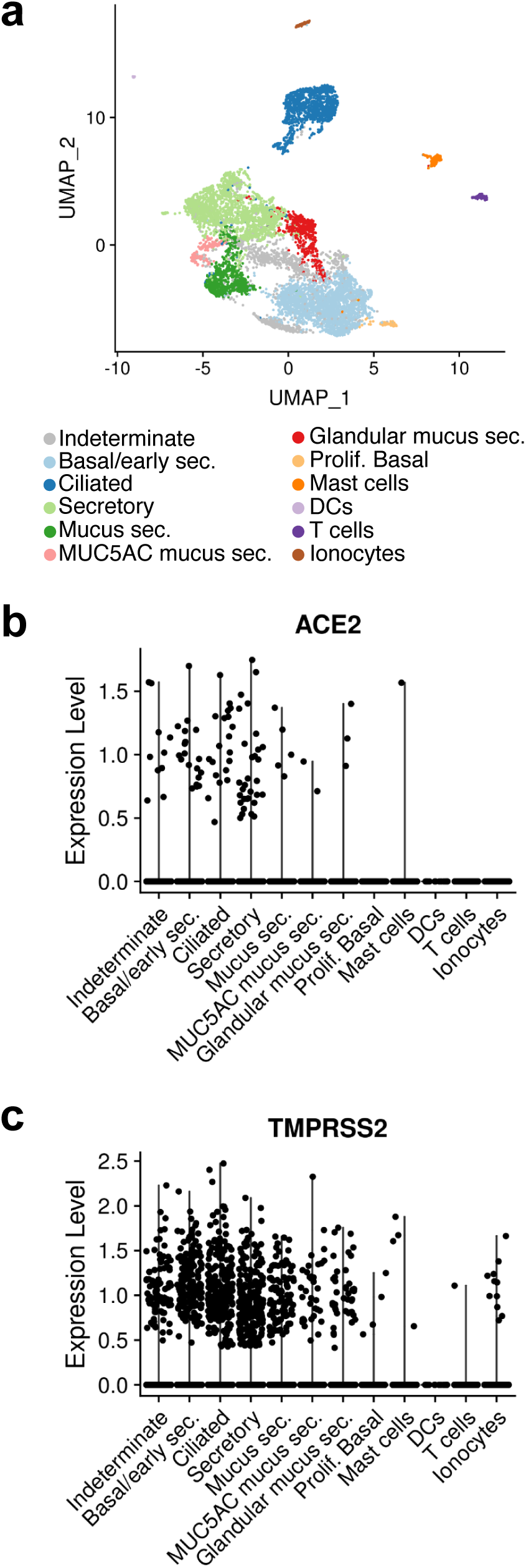
*ACE2* and *TMPRSS2* are expressed by multiple nasal airway epithelial cell types. (a) UMAP visualization of cells derived from a human nasal airway epithelial brushing depicts multiple epithelial and immune cell types identified through unsupervised clustering. (b) Normalized expression of *ACE2* in epithelial and immune cell types. (c) Normalized expression of *TMPRSS2* in epithelial and immune cell types.

### *TMPRSS2* is part of a mucus secretory co-expression network highly induced by T2 inflammation

We next sought to determine the variation in nasal epithelial expression of *ACE2* and *TMPRSS2* across healthy and asthmatic children, and to identify biological mechanisms that regulate this variation. Thus, we performed weighted gene co-expression network analysis (WGCNA) on whole transcriptome sequencing data from nasal airway brushings of 695 Puerto Rican healthy and asthmatic children in the Genes-Environments and Asthma in Latino Americans II study (GALA II). This analysis identified 54 co-expression networks representing cell type-specific expression programs such as ciliogenesis, mucus secretion, and pathways of immunity and airway inflammation (Supplementary Table 2). The *TMPRSS2* gene was contained within one of a set of three highly correlated networks exhibiting strong enrichments for mucus secretory cell genes and pathways (Figure 2a, Supplementary Table 2,3). For example, the black network, which was highly correlated with *TMPRSS2* expression (r=0.64, p=1e-82), was strongly enriched for *Golgi mediated transport* and *COPI-dependent Golgi to ER transport* pathways, both of which are involved in the normal processing and transport of mucin proteins (Figure 2a). *TMPRSS2* itself fell within and was highly correlated with expression of the pink network (r=0.68, p=3e-97), which was highly enriched for *mucus goblet cell* markers (p=2e-6, Figure 2a,b). The pink network was also enriched for genes involved in the *O-linked glycosylation of mucins* pathway (p=9e-4), which is vital to the function of mucus secretory cells, especially those induced by T2 inflammation (r=0.68, p=3e-97, Figure 2a,b). In fact, we found that this network contained the T2 cytokine *IL13* while being particularly enriched for genes known to mark and transcriptionally regulate IL-13-induced mucus metaplasia (*FCGBP, SPDEF, FOXA3*). The saddle brown network was also related to mucus secretory cells, and contained the most canonical T2 inflammation markers^11, 23^ including *POSTN, CLCA1, CPA3, IL1RL1, CCL26*, and was strongly correlated with both *TMPRSS2* (r=0.61, p=5e-72, Figure 2c) and the other T2 mucus secretory network (pink) (r=0.92, p=3e-280, Supplementary Table 4). In contrast, we found *ACE2* expression to be strongly negatively correlated with expression of both T2 networks (pink: r=-0.61, p=3e-72, saddle brown: r=-0.7, p=2e-102, Figure 2e,f). To identify subjects with high and low T2 inflammation, we hierarchically clustered all subjects based on the expression of genes in the canonical T2 network (saddle brown). This resulted in the identification of two distinct groups we labeled as T2-high (n=364) and T2-low (n=331) (Supplementary Figure 1a). We found that this expression-derived T2 status was strongly associated with traits known to be driven by T2 inflammation including IgE levels, exhaled nitric oxide (FeNO), blood eosinophils, and asthma diagnosis (Supplementary Figure 1b-e). Notably, *TMPRSS2* levels were 1.3-fold higher in T2-high subjects (p=1e-62), while, *ACE2* expression was 1.4-fold lower in T2-high subjects (p=2e-48) (Figure 2d,g).

**Figure 2.**
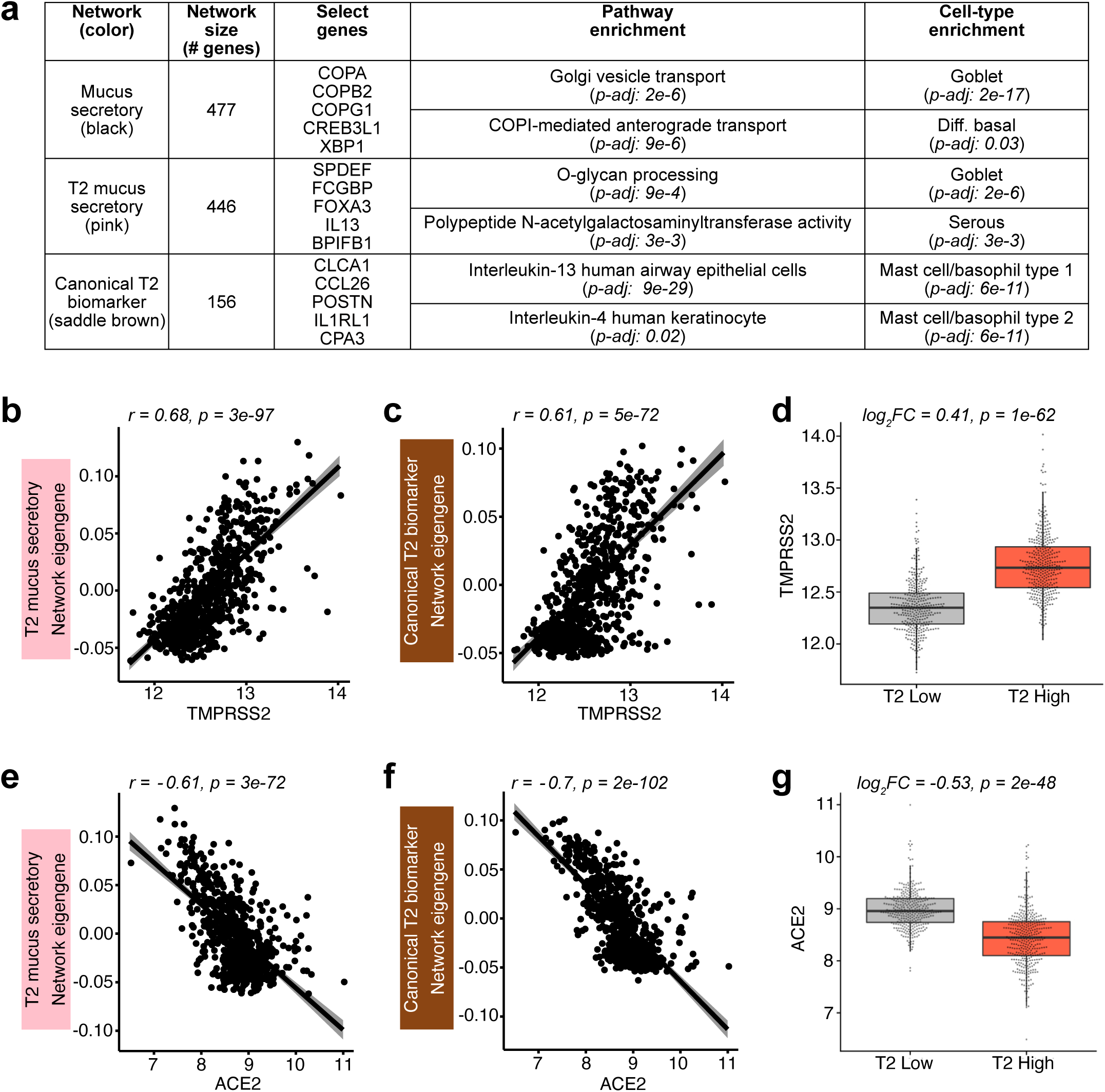
*TMPRSS2* is a mucus secretory network gene regulated by T2 inflammation. (a) WGCNA identified networks of co-regulated genes related to mucus secretory function (black), T2 inflammation-induced mucus secretory function (pink), and canonical T2 inflammation biomarkers (saddle brown). *TMPRSS2* was within the pink network. Select pathway and cell type enrichments for network genes are shown. (b) Scatterplot revealing a strong positive correlation between *TMPRSS2* expression and summary (eigengene) expression of the T2 inflammatory, mucus secretory network. (c) Scatterplot revealing a strong positive correlation between *TMPRSS2* expression and summary (eigengene) expression of the canonical T2 inflammation biomarker network. (d) Box plots revealing strong upregulation of *TMPRSS2* expression among T2-high compared to T2-low subjects. (e) Scatterplot revealing a strong negative correlation between *ACE2* expression and summary (eigengene) expression of the T2 inflammation mucus secretory network. (f) Scatterplot revealing a strong negative correlation between *ACE2* expression and summary (eigengene) expression of the canonical T2 inflammation biomarker network. (g) Box plots revealing strong downregulation of *ACE2* expression among T2-high compared to T2-low subjects.

To investigate whether the strong *in vivo* relationship between airway T2 inflammation and *TMPRSS2*/*ACE2* expression is causal in nature, we performed *in vitro* stimulation of paired air-liquid interface (ALI) mucociliary airway epithelial cultures with 72 hours of IL-13 or mock stimulus (n=5 donors, Figure 3a). Performing paired differential expression analysis between the mock and IL-13 stimulated cultures, we found that *ACE2* and *TMPRSS2* were strongly down- and up-regulated, respectively, supporting our *in vivo* analysis results (log_2_FC= -0.67, p=5e-3, log_2_FC= 1.20, p=5e-9, Figure 3b,c). To better understand the cellular basis of *TMPRSS2* and *ACE2* regulation by IL-13, we leveraged scRNA-seq data previously generated on tracheal airway epithelial cultures that were chronically stimulated (10 days) with IL-13 or control media (Figure 3a,d). Similar to our results from *in vivo* nasal scRNA-seq data, we observed that *ACE2* expression was highest among basal, ciliated, and early/intermediate secretory cell populations, with *ACE2* being significantly downregulated by IL-13 among both basal and intermediate secretory cells (Figure 3e). Also mirroring the *in vivo* scRNA-seq data, *TMPRSS2* was expressed across all epithelial cell types, but at a higher frequency among secretory cells (Figure 3f). IL-13 stimulation induced dramatic upregulation of *TMPRSS2* in early secretory, intermediate secretory, and mature mucus secretory cell populations (Figure 3f). Furthermore, IL-13 stimulated mucus metaplasia that resulted in the development of a novel mucus secretory cell type and an IL-13 inflammatory epithelial cell that both highly expressed *TMPRSS2* (Figure 3f). Together, our *in vivo* and *in vitro* analyses strongly suggest that *TMPRSS2* is part of a mucus secretory cell network that is highly induced by IL-13-mediated T2 inflammation.

**Figure 3.**
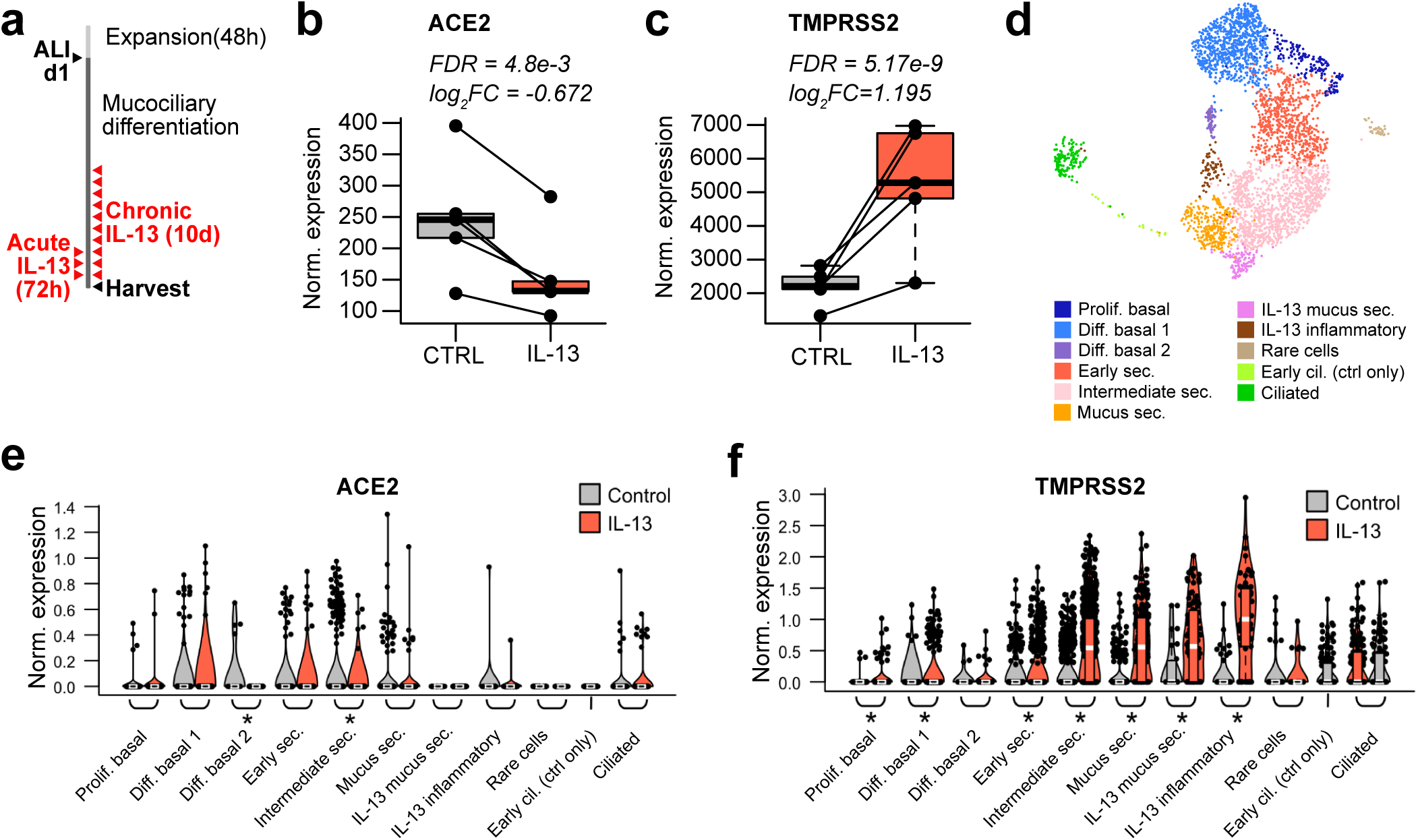
*ACE2* and *TMPRSS2* expression are both regulated by IL-13 in the mucociliary airway epithelium. (a) Experimental schematic detailing the timeline for differentiation of basal airway epithelial cells into a mucociliary airway epithelium and treatment with chronic (10 days) or acute (72 hours) IL-13 (10ng/ml). (b) Box plots of count-normalized expression between paired nasal airway cultures (control/IL-13) revealing strong downregulation of bulk *ACE2* expression with IL-13 treatment. Differential expression results are from DESeq2. (c) Box plots of count-normalized expression between paired nasal airway cultures (control/IL-13) revealing strong upregulation of bulk *TMPRSS2* expression with IL-13 treatment. Differential expression results are from DESeq2. (d) UMAP visualization of cells derived from control and IL-13 stimulated tracheal airway ALI cultures depict multiple epithelial cell types identified through unsupervised clustering. (e) Violin plots of normalized *ACE2* expression across epithelial cell types from tracheal airway ALI cultures, stratified by treatment (gray = control, red = IL-13). Differential expression using a Wilcoxon test was performed between control and IL-13-stimulated cells with significant differences in expression for a cell type indicated by a * (p < 0.05). (f) Violin plots of normalized *TMPRSS2* expression across epithelial cell types from tracheal airway ALI cultures, stratified by treatment (gray = control, red = IL-13). Differential expression using a Wilcoxon test was performed between control and IL-13-stimulated cells with significant differences in expression for a cell type indicated by a * (p < 0.05).

### *ACE2* belongs to an interferon response network that is induced by respiratory virus infections

Returning to the *in vivo* nasal airway epithelial expression networks, we found that *ACE2* expression was highly correlated with expression of two networks (purple and tan) (purple: r=0.74, p=3e-120, tan: r=0.72, p=2e-110, Figure 4a,b). The purple network was highly enriched for genes that mark cytotoxic T cells and antigen-presenting dendritic cells, both of which are particularly abundant in a virally infected epithelium (Figure 4c, Supplementary Table 2), whereas the tan network was strongly enriched for interferon and other epithelial viral response genes (*IFI6, IRF7, CXCL10, CXCL11*) (Figure 4c, Supplementary Table 2). Clustering of subjects based on the interferon response network genes resulted in two groups, one highly (interferon-high=78) and one lowly (interferon-low=617) expressing these interferon response network genes (Supplementary Figure 2). We found that *ACE2* expression was 1.7-fold higher in the interferon-high vs. interferon-low group (Figure 4d). In a previous study, we found that children with nasal gene expression characteristic of the interferon network tended to be infected with a respiratory virus, despite being asymptomatic^19^. To explore the possibility of this relationship in our current dataset, we metagenomically analyzed the RNA-seq data for all subjects to identify those harboring reads for a respiratory virus. This analysis found that 18% of GALA II children were asymptomatically harboring a respiratory virus from one of eight general respiratory virus groups (Figure 4e). Strikingly, we found that 78% of interferon-high subjects were virus carriers compared to only 10% of interferon-low subjects. These results demonstrate how asymptomatic virus carriage nonetheless stimulates an active viral response that includes *ACE2*.

**Figure 4.**
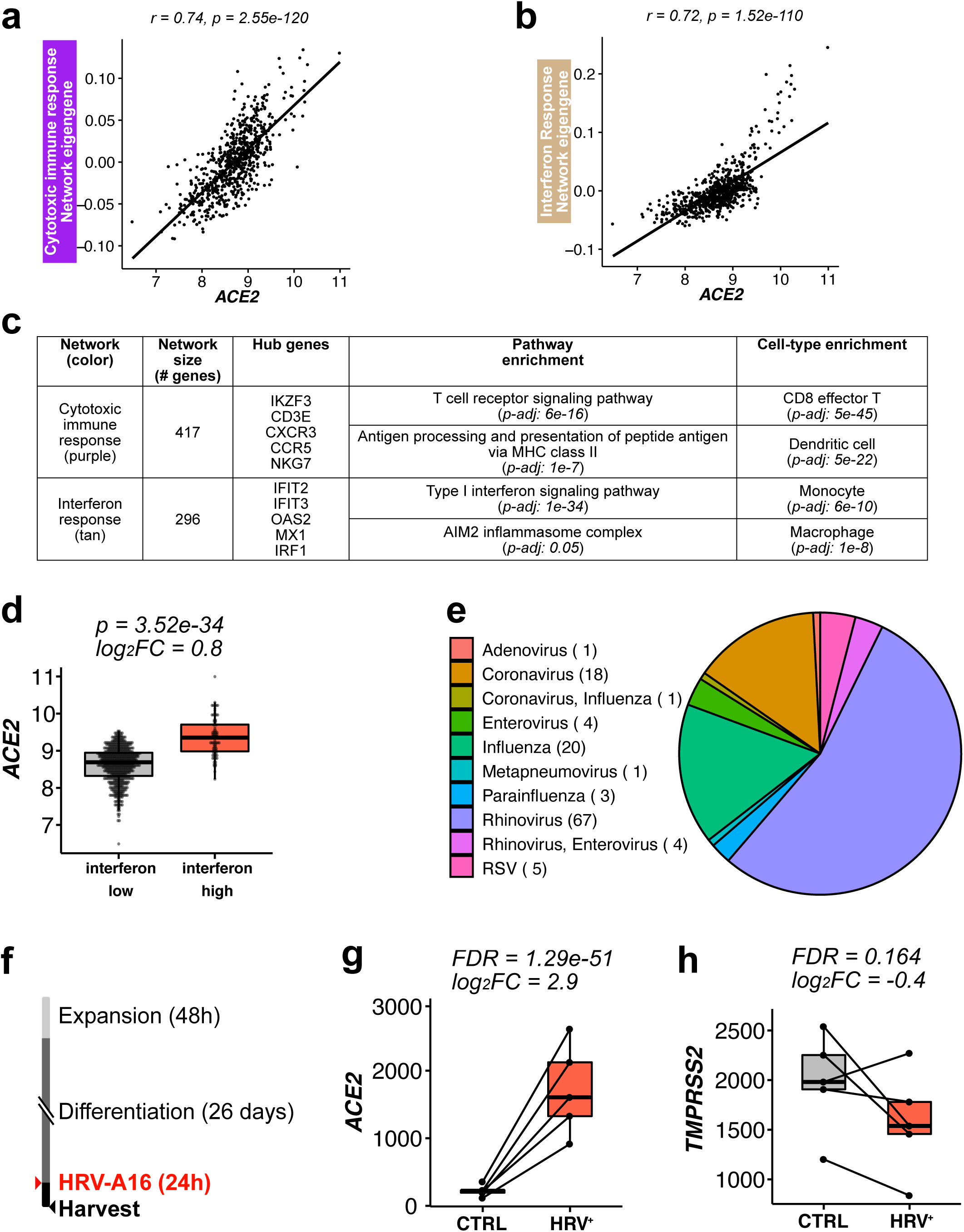
*ACE2* is an interferon response network gene regulated by respiratory virus infections. (a) Scatter plot revealing a strong positive correlation between *ACE2* expression and summary (eigengene) expression of the cytotoxic immune response network (purple). (b) Scatterplot revealing a strong positive correlation between *ACE2* expression and summary (eigengene) expression of the interferon response network (tan). (c) WGCNA analysis identified networks of co-regulated genes related to cytotoxic immune response (purple) and interferon response (tan). *ACE2* was within the purple network. Select pathway and cell type enrichments for network genes are shown. (d) Box plots of count-normalized expression from GALA II nasal epithelial samples reveal strong upregulation of *ACE2* expression among interferon-high compared to interferon-low subjects. Differential expression results are from DESeq2. (e) Pie graph depicting the percentage of each type of respiratory virus infection found among GALA II subjects in whom viral reads were found. (f) Experimental schematic detailing timeline for differentiation of basal airway epithelial cells into a mucociliary airway epithelium and experimental infection with HRV-A16. (g) Box plots of count-normalized expression between paired nasal airway cultures (control/HRV-A16 infected) revealing strong upregulation of *ACE2* expression with HRV-A16 infection. Differential expression results are from DESeq2. (h) Box plots of count-normalized expression between paired nasal airway cultures (control/HRV-A16-infected) revealing no effect of HRVA-16 on *TMPRSS2* expression. Differential expression results are from DESeq2.

To directly test the effect of respiratory virus infection on epithelial *ACE2* gene expression we again employed our ALI mucociliary epithelial culture system. Performing mock or human rhinovirus-A16 infection of mature cultures (Day 27, Figure 4f) from 5 donors we found 7.7-fold upregulation of *ACE2* gene expression with HRV-A infection (p=1.3e-51, Figure 4g). In contrast, we only observed a trend for down regulation of *TMPRSS2* gene expression among virally infected subjects (Figure 4h). These results confirm the strong regulation of *ACE2* gene expression by viral infection.

### Genetic determinants of *ACE2* and *TMPRSS2* expression in the nasal airway epithelium

We next explored the role of genetic regulatory variants in helping to drive epithelial expression of *ACE2* and *TMPRSS2.* To do this, we performed cis-eQTL analysis for these two genes, using nasal gene expression and genome-wide genetic variation data collected from the GALA II study children. We identified 316 and 36 genetic variants significantly associated with expression of *ACE2* and *TMPRSS2*, respectively (Figure 5a,b). Stepwise forward-backward regression analysis of these eQTL variants revealed a single independent eQTL variant (rs181603331) for the *ACE2* gene (6e-23), located ∼20kb downstream of the transcription start site (Figure 5a). This rare eQTL variant (allele frequency [AF]=1%) was associated with a large decrease in *ACE2* expression (log_2_A_FC_=-1.6) (Figure 5c).

**Figure 5.**
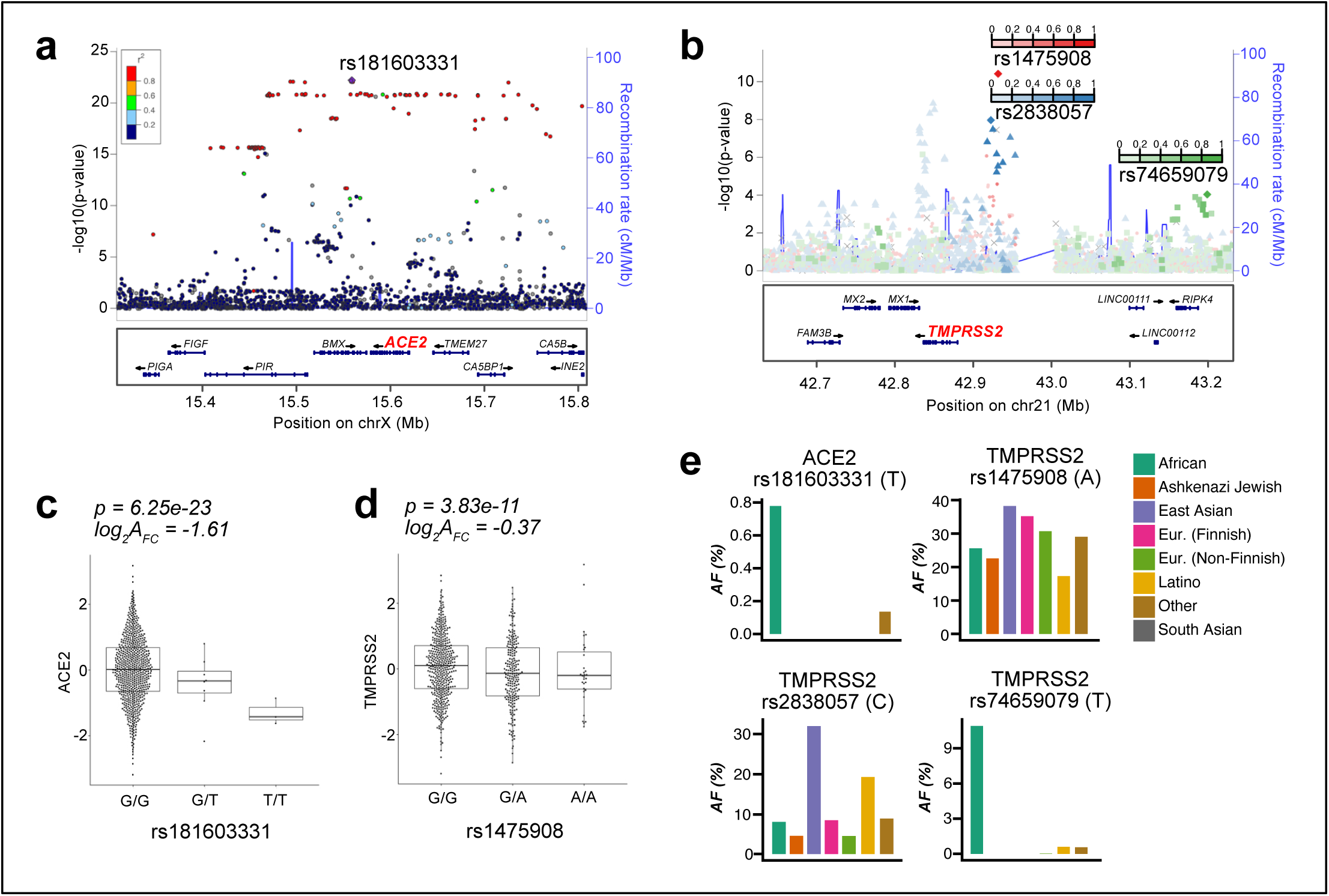
*ACE2* and *TMPRSS2* nasal airway expression are regulated by eQTL variants. (a) Locuszoom plot of *ACE2* eQTL signals. The lead eQTL variant (rs18160331) is highlighted with a purple dot. The strength of Linkage Disequilibrium (LD) between rs18160331 and other variants is discretely divided into five quantiles and mapped into five colors (dark blue, sky blue, green, orange, and red) sequentially from low LD to high LD. (b) Locuszoom plot of *TMPRSS2* eQTL signals. The three independent eQTL variants (rs1475908, rs2838057, rs74659079) and their LD with other variants (r^2^) are represented by red, blue, and green color gradient respectively. (c) Box plots of normalized *ACE2* expression among the three genotypes of the lead *ACE2* eQTL variant (rs18160331). log_2_A_FC_ = log2 of the allelic fold change associated with the variant. (d) Box plots of normalized *TMPRSS2* expression among the three genotypes of the lead *TMPRSS2* eQTL variant (rs1475908). log_2_A_FC_ = log2 of the allelic fold change associated with the variant. (e) Bar plots depicting allele frequencies of the *ACE2* eQTL variant rs18160331 and *TMPRSS2* eQTL variants (rs1475908, rs2838057, rs74659079) across world populations. Allele frequency data were obtained from gnomAD v2.1.1.

Similar analysis of the *TMPRSS2* eQTL variants yielded three independent eQTL variants (rs1475908 AF=20%, rs74659079 AF=4%, and rs2838057 AF=13%, Figure 5b). The eQTL variant rs1475908 was associated with a decrease in *TMPRSS2* expression (log_2_A_FC_=-0.37, Figure 5d), whereas both the rs74659079 and rs2838057 eQTL variants were associated with increased *TMPRSS2* expression (log_2_A_FC_=0.38, 0.43, respectively, Supplementary Figure 3).

Examining the frequency of these eQTL variants among eight world populations listed in the gnomAD genetic variation database (v2.1.1), we found that the *ACE2* eQTL variant was only present in people of African descent and at a low frequency (AF=0.7%, Figure 5e). In contrast, the *TMPRSS2* eQTL variant associated with decreased expression, rs1475908, occurred across all world populations, with the highest allele frequencies among East Asians (AF=38%), Europeans (AF=35%), intermediate frequencies among Africans (AF=26%) and Ashkenazi Jews (AF=23%), and the lowest frequency among Latinos (AF=17%). The two *TMPRSS2* eQTL variants associated with increased expression exhibited much more disparate allele frequencies across world populations. Namely, the allele frequency of rs74659079 is above 1% only among people of African descent (AF=11%) and 4% in the participating Puerto Rican population. Likewise, the rs2838057 eQTL variant, which was associated with increased *TMPRSS2* expression was present at a frequency of 32% in East Asians, 20% in Latinos, and <10% in all other world populations. Together, these results suggest that if *TMPRSS2* levels influence susceptibility to SARS-CoV-2, then genetics may play a significant role in infection risk and that this risk will vary significantly across world populations.

### Multi-variable modeling of airway *ACE2* and *TMPRSS2* gene expression

Our analyses indicate that T2 inflammation, interferon/viral response signaling, and genetics are all determinants of *ACE2* and *TMPRSS2* gene expression in the airway epithelium of children. Therefore, we next sought to determine the relative importance of these factors in determining levels of these genes using multi-variable regression analysis. We included asthma status, age, and sex as model covariates since chronic lung disease, increasing age, and male sex have all been associated with increased risk of poor COVID-19 illness outcomes. Modeling *ACE2* expression among GALA II children, we found that T2 and interferon statuses had the strongest effects on *ACE2* expression (p=1.6e-57, p=6.5e-43, respectively), with T2-low and interferon-high individuals exhibiting the highest levels of expression. These two variables independently explained 24% and 17% of the variance in *ACE2* expression (Table 1). While the *ACE2* eQTL variant, rs181603331, was associated with a notable decrease in *ACE2* levels, it only accounted for 1.2% of the variance, reflecting the low frequency of this variant in our population. Increasing age and asthma diagnosis were both associated with small decreases in *ACE2* expression, although both variables accounted for less than 2% of the variance, and sex was not a significant predictor (Table 1).

Similar modeling of *TMPRSS2* expression found that T2-high status dramatically increased expression, with an effect size 5.4x larger than any other variable, capturing 33% of total variation in *TMPRSS2* (Table 1). While statistically significant, the two *TMPRSS2* eQTL variants associated with increased expression exhibited small effect sizes totaling <1% of variance explained. All other predictors were not significant. Collectively, these modeling results confirm that both T2 and interferon inflammation are strong and antagonistic regulators of *ACE2* expression and show that T2 inflammation is the lone dominant driver of airway expression of *TMPRSS2.*

### Coronavirus Infections drive an enhanced cytotoxic immune response

Our metagenomic analysis of RNA-seq data from the nasal brushings identified 18 children with viral sequence reads from one of four different coronavirus (CoV) species (OC43, JKU1, 229E, NL63) (Supplementary Table 5). This allowed us to explore airway transcriptomic responses to infection with coronavirus subfamily viruses specifically, which will likely most resemble responses to SARS-CoV-2. To increase the likelihood that these subjects were experiencing an active viral infection, we limited our analysis to the 11 most highly infected subjects, comparing them to all subjects not infected with a virus (n=571). To allow us to discriminate CoV-enhanced responses from those that are more general to respiratory viruses, we also established a virus control group composed of the 37 subjects highly infected with human rhinovirus species (HRV) (Supplementary Table 6). We first compared expression of genes in the cytotoxic immune response (purple) network and interferon response (tan) network (discussed earlier; see Figure 4a, b) among these virus infected groups, and found that both networks were more highly expressed in virus-infected individuals (Figure 6a, b). Moreover, while the induction in interferon response was similar for both CoV and HRV groups, induction in the cytotoxic immune response was considerably higher in CoV-infected (ΔE_g_ = 0.049) compared to HRV-infected individuals (ΔE_g_ = 0.032, Figure 6b).

**Figure 6.**
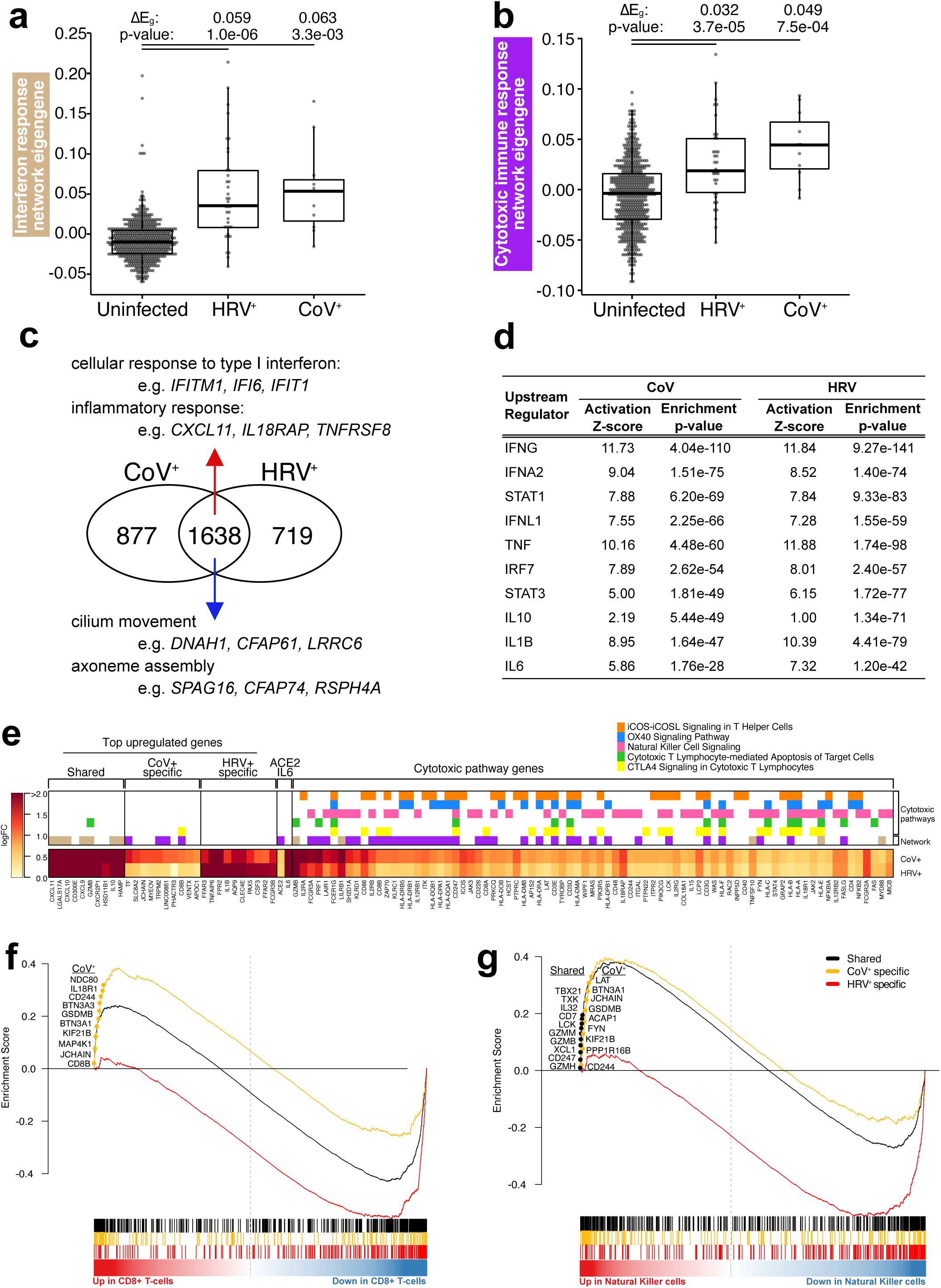
Coronavirus infections elicit an enhanced cytotoxic immune response from the airway epithelium. (a) Box plots revealing a strong and equivalent upregulation of summary (eigengene [E_g_]) expression for the interferon response network among HRV and CoV-infected GALA II subjects, compared to uninfected subjects. (b) Box plots revealing upregulation in summary (eigengene) expression for the cytotoxic immune response network among HRV-infected GALA II subjects that is even stronger for the CoV infected group. (c) Venn Diagram describing the number of differentially expressed genes in HRV and CoV infected groups compared to the uninfected group, and the extent of their overlap. For genes differentially expressed in both groups, select enriched pathways and underlying genes that are highly differentially expressed are shown. (d) Top upstream regulators predicted by Ingenuity Pathway Analysis to be regulating the genes that were upregulated in CoV. Enrichment values for these CoV regulators, using the HRV upregulated genes are also shown. (e) Heatmap of the log_2_FC in gene expression for CoV and HRV groups when compared to the uninfected group. Top significantly upregulated genes are shown, along with *ACE2, IL6*, and genes identified as belonging to cytotoxic pathways, which were enriched within the virally upregulated CoV group DEGs based on IPA canonical pathway analysis. Color bars indicate which WGCNA network and or IPA canonical pathway each gene belongs to. (f) Gene set enrichment analysis plot for CD8+ T cells. The black (shared), yellow (CoV-enhanced), and red (HRV-enhanced) curves display the enrichment score for the indicated viral gene set as the analysis walks down the ranked distribution of genes ordered by fold change in expression between CD8+ T cells relative to all other immune cell types (red-blue color bar). Genes are represented by vertical bars in the same color as the curve of the viral gene group they represent. Denoted genes are a representative set from the leading edge (most responsible for the enrichment). (g) Gene set enrichment analysis plot for NK cells. The black (shared), yellow (CoV-enhanced), and red (HRV-enhanced) curves display the enrichment score for the indicated viral gene set as the analysis walks down the ranked distribution of genes ordered by fold change in expression between NK cells relative to all other immune cell types (red-blue color bar). Genes are represented by vertical bars in the same color as the curve of the viral gene group they represent. Denoted genes are from the leading edge (most responsible for the enrichment).

To further explore this increase in cytotoxic immune response and other potential pathways in CoV-infected individuals, we next performed a transcriptome-wide screen for genes differentially expressed in CoV or HRV-infected groups compared to uninfected individuals. These analyses revealed 2,515 differentially expressed genes (DEGs) with CoV infection and 2,357 DEGs with HRV infection (FDR < 0.05 and log_2_FC > |0.5|), of which 35% and 31% were only observed with CoV and HRV infections, respectively, based on our significance cutoff (Figure 6c). Upstream regulator analysis with IPA carried out separately on CoV and HRV infection response genes showed that the top cytokines and transcription factors that may regulate these infections are shared between the two virus families, including IL10, IL1B, IFNG, IFNA2, and STAT1 (Figure 6d). One inferred upstream regulator of CoV response genes, IL-6, which was also among the genes upregulated with CoV infection (log_2_FC=2.2, Figure 6e), is especially noteworthy considering that an IL-6 blocking antibody therapy is currently under investigation for use in treatment of COVID-19 illnesses^24^. Additionally, we found *ACE2* among the shared upregulated genes, reinforcing its upregulation in the course of different respiratory virus infections (log_2_FC in CoV^+^=0.6, log_2_FC in HRV^+^=0.5, Figure 6e).

In trying to understand the biological basis of the viral responses we found to be CoV-specific in our differential expression analysis, we considered whether the differential presence and/or response of various immune cell types was an explanatory factor. To investigate this, we used gene set enrichment analysis (GSEA) to test for enrichment of CoV-specific, HRV-specific, and CoV/HRV-shared DEG sets among gene markers for 11 different flow-sorted human immune cell types defined based on whole transcriptome data (citation) (Supplementary Table 7). The shared viral DEGs showed significant enrichment for genes characteristic of macrophages, monocytes, neutrophils, dendritic cells, and NK cells. In contrast, the set of CoV-enhanced DEGs resulted in strong enrichments for both CD8+ T cells and dendritic cells, suggesting an especially important role for activation of cytotoxic T cells though antigen presentation by dendritic cells in CoV infections (Figure 6f). Also supporting an enriched cytotoxic response among CoV-infected subjects was a strong enrichment for CoV-specific DEGs among NK cells, which participate heavily in the killing of virally infected cells (Figure 6g). We note that these enrichments were not observed among HRV-enhanced DEGs, which were instead most strongly enriched among neutrophils, as well as eosinophils, macrophages, and monocytes. Furthermore, through pathway analysis we identified multiple pathways related to cytotoxic T cell and NK cell activity that were enriched either specifically or more dramatically among CoV DEGs compared to HRV DEGs (Figure 6e). These results suggest that while CoV infections are highly similar to HRV infections, they likely elicit an enhanced cytotoxic immune response.

## Discussion

Although the high variability in clinical outcomes of COVID-19 illness is now well documented and multiple demographic and clinical traits have been associated with severe disease, little is known about the host biologic factors underlying this variability. In the current study, we reasoned that population variation in upper airway expression of the ACE2 receptor for SARS-CoV-2 and the virus-activating TMPRSS2 protease, would drive infection susceptibility and disease severity. We therefore deployed network and eQTL analysis of nasal airway epithelial transcriptome data from a large cohort of healthy and asthmatic children to determine mechanisms associated with airway expression of these genes, and their relative power in explaining variation in the expression of these genes among children. We observed only weak associations with asthma status, age, and gender among children aged 8-21 years. Moreover, although we found that genetics does influence expression of these genes, the effect of this variation was small in comparison to the dramatic influence of T2 cytokine-driven inflammation on both *ACE2* (downregulation) and *TMPRSS2* (upregulation) expression levels. We found an equally important role for viral-driven interferon inflammation in regulating levels of *ACE2* in the airway. Additionally, through study of *in vivo* upper airway CoV subfamily infections, we not only identify inflammatory regulators of these infections, but also provide evidence that this subfamily of viruses drives an enhanced cytotoxic immune response. Our work provides a set of biomarkers that can be easily examined in COVID-19 patients, through analysis of nasal swabs, to determine the relative importance of these mechanisms and genes in governing susceptibility to infection, severe illness and death.

Our single cell analysis of an *in vivo* nasal brushing observed *ACE2* expression, albeit at low frequency, primarily among basal, ciliated, and less mature, early secretory cells. These results are supported by a recent report of *ACE2* expression in transient secretory cells, likely a close equivalent to our early secretory population^25^. Although a much higher portion of cells, representing all epithelial cell types, expressed *TMPRSS2*, the low frequency of *ACE2*^+^ cells resulted in very few dual *ACE2*/*TMPRSS2* expressing cells. However, we caution that a cell may not need to be *TMPRSS2*^+^ to be susceptible to infection, since it has been demonstrated the TMPRSS2 protein is secreted from nasal airway epithelial cells^26^. We also caution that scRNA-seq data are known to exhibit biases in gene detection, and thus the level and frequency of *ACE2* expression across cells may be much higher than we observe here. In line with this possibility we observe more moderate levels of *ACE2* expression in our bulk RNA-seq data on nasal brushings.

Airway inflammation caused by type 2 cytokine production from infiltrating immune cells plays a prominent role in the control of cellular composition, expression, and thus biology of the airway epithelium^11, 13, 23, 27^. Moreover, while T2 airway inflammation is an important driver of T2-high asthma and COPD disease endotypes, it is also associated with atopy in the absence of lung disease, a very common phenotype in both children and adults. In fact, among the children in this study, we find that 43% of non-asthmatics were scored as T2-high based on expression profile, further substantiating the high prevalence of T2 airway inflammation outside of those with lung disease. Our data suggest that airway epithelial *TMPRSS2* expression is highly upregulated by T2 inflammation, and specifically by IL-13. Both our network and single cell data show that *TMPRSS2* is most prominent in less developed “early secretory” cells as well as in more mature mucus secretory cells. Based on our *in vitro* data, IL-13 upregulates *TMPRSS2* across nearly all types of epithelial cells, but the core of this effect appears to be in the metaplastic mucus secretory cells that are generated as a consequence of IL-13 signaling^14, 15^. In fact, our network data suggest that, although *TMPRSS2* expression is highly correlated with that of a co-expressed network of mucus secretory genes characterizing “normal”, non-metaplastic, mucus secretory cells, it’s correlated even more strongly with a network that characterizes mucus secretory cells undergoing IL-13-induced metaplasia. In contrast to enhanced levels of *TMPRSS2*, T2 inflammation, whether observed *in vivo* or induced with IL-13 stimulation, precipitated a dramatic reduction in levels of epithelial *ACE2*, thus complicating expectations for how T2 inflammation might affect overall risk for a poor COVID-19 outcome. Germane to this question, a recent study of 85 fatal COVID-19 subjects found that 81.2% of them exhibited very low levels of blood eosinophil levels^4^. Blood eosinophil levels are a strong, well-known predictor of airway T2 inflammation and were strongly correlated with T2 status in our study as well^11, 23^. Together, these studies provisionally suggest that T2 inflammation may predispose individuals to experience better COVID-19 outcomes through a decrease in airway levels of *ACE2* that override any countervailing effect from increased expression of *TMPRSS2*. However, both *in vitro* experiments examining IL-13 effects on SARS-CoV-2 infection and empirical data on COVID-19 outcomes among T2-high and T2-low patients will certainly be needed to determine whether this common airway inflammatory endotype ultimately protects against or exacerbates COVID-19 illness. As mentioned above, we note that measurement of blood eosinophil levels could be used as an informative and more accessible (albeit less powerful) proxy for investigating the association between airway T2 inflammation and outcomes of COVID-19. Moreover, given the higher frequency of T2 inflammation among asthmatic subjects, this population should be monitored especially closely given the enhanced risk of complications due to respiratory virus infection in those with asthma.

In addition to a strong negative influence of T2 inflammation on *ACE2* expression in the airway, we found an equally strong positive influence of respiratory virus infections on levels of this gene. Network analysis placed *ACE2* within an interferon viral response network suggesting that these cytokines are a driving force behind *ACE2* upregulation. This information is interesting in several regards. First, it suggests that SARS-CoV-2 and other coronaviruses using ACE2 as a receptor could leverage the host anti-viral response to increase the infectability of airway cells. Secondly, as data here and elsewhere show, asymptomatic carriage of respiratory viruses is common, especially in young children^19, 28-31^. Children in the GALA II cohort included in this study ranged in age from 8-21 years; among them we found 18% who were carrying respiratory viruses without illness. However, as we show in this and our previous study^19^, even asymptomatic carriage of respiratory viruses exacts a fundamental change in airway epithelial expression and immune cell presence, including upregulation of *ACE2* expression. In determining outcomes, this potential detrimental influence of virus carriage may also be weighed against a potentially beneficial influence of virus carriage through a more potent cross serologic-immune defense in these individuals, especially if the virus carried is a coronavirus family member. Ultimately, the effect of current or recent virus carriage on COVID-19 outcomes will need to be determined by *in vivo* studies in patients, followed up with controlled *in vitro* studies of virally infected cells. At any rate, the apparent dependence of *ACE2* expression on interferon signaling suggests that targeted blockade of this interferon effect could control SARS-CoV-2 infection.

Our evaluation of genetic influences on airway *ACE2* and *TMPRSS2* expression revealed a single rare eQTL for *ACE2* and several more frequent eQTL variants for *TMPRSS2*. While both the effect size and explanatory power of these variants paled in comparison to the influence of T2 inflammation and interferon signaling in multi-variable modeling of expression for these genes, the effect of these variants may still be strong enough to alter infection rates and or illness severity, especially in the populations where these variants are most frequent. Thus, future genetic studies of COVID-19 should pay particular attention to these eQTL variants.

A particularly vexing question regards the mechanisms that underlie the unusual severity of illness associated with SARS-CoV-2, especially when compared to most circulating respiratory viruses. Clearly, severe disease often entails development of pneumonia, possibly resulting from an expanded tropism of SARS-CoV-2 to include lower lung airway and alveolar cells. The most severe patients also appear to experience an exuberant immune response, characterized a “cytokine storm”^24^, occurring with and possibly driving the development of acute respiratory distress syndrome (ARDS). Supposing that aspects of epithelial response to coronavirus family members would be shared, including with SARS-CoV-2, we examined *in vivo* coronavirus infection among the GALA II children. We found that CoV infections elicit a broad airway transcriptome response, similar to HRV infections, and we identified a panel of cytokines and transcription factors that likely regulate these responses. In particular, we found that IL-6 was predicted to regulate responses to CoV and was itself upregulated with these infections. These data support the recent investigation of tocilizumab (IL-6R blocking antibody) for the treatment of COVID-19 illnesses^24^. Strikingly our analysis revealed an increased cytotoxic immune response with CoV infection, driven by the differential presence and activity of cytotoxic CD8+ T cells and NK cells, as compared to the more heavily neutrophil-based responses to HRV infection. Although preliminary, this finding, if similarly occurring with SARS-CoV-2 infection, could partly explain the dramatic inflammation observed in SARS-CoV-2 patients, which can extend to the distal lung.

In summary, our data suggests that the strongest determinants of airway *ACE2* and *TMPRSS2* expression are T2 inflammation and viral-induced interferon inflammation, with limited but noteworthy influence from genetic variation. Whether these factors drive better or worse clinical outcomes remains to be determined, but closely watching individuals with these airway endotypes in the clinical management of COVID-19 illnesses would be prudent.

## Methods

### MATERIALS AND CORRESPONDENCE

Further information and requests for resources and reagents should be directed to and will be fulfilled by Max A. Seibold, Ph.D.

### EXPERIMENTAL METHODS

#### Human subject information

Under the Institutional Review Board (IRB) approved Asthma Characterization Protocol (ACP) at National Jewish Health (HS-3101) we consented a 56 year old asthmatic subject, from which we collected nasal airway epithelial cells. The nasal airway cells were brushed from the inferior turbinate using a cytology brush and used for the scRNA-seq experiment described in Figure 1. Nasal airway epithelial cells used for bulk RNA-seq network and eQTL analysis came from GALA II study subjects described below. Nasal airway epithelial cell ALI culture experiments all used cells derived from GALA II study subjects. Human tracheal airway epithelial cells used for *in vitro* IL13 stimulation and scRNA-seq experiment were isolated from a single de-identified lung donor obtained from the International Institute for the Advancement of Medicine (Edison, NJ), and Donor Alliance of Colorado. The National Jewish Health Institutional Review Board (IRB) approved our research on the tracheal airway epithelial cells under IRB protocol HS-3209. These cells were processed and given to us through the National Jewish Health (NJH) live cell core, which is an institutional review board-approved study (HS-2240) for the collection of tissues from consented patients for researchers at NJH.

#### GALA II study subjects

The Genes-Environment & Admixture in Latino Americans study (GALA II) is an on-going case-control study of asthma in Latino children and adolescents. GALA II was approved by local institutional review boards (UCSF, IRB number 10–00889, Reference number 153543, NJH HS-2627) and all subjects and legal guardians provided written informed assent and written informed consent, respectively^32, 33^. A full description of the study design and recruitment has been previously described elsewhere^32-34^. Briefly, the study includes subjects with asthma and healthy controls of Latino descent between the ages of 8 and 21, recruited from the community centers and clinics in the mainland U.S. and Puerto Rico (2006-present). Asthma case status was physician-diagnosed. Recruited subjects completed in-person questionnaires detailing medical, environmental, and demographic information. Physical measurements including spirometry were obtained, and subjects provided a blood sample for DNA extraction and later Whole Genome Sequencing. GALA subjects that were part of this analysis were all recruited from Puerto Rico (n=695). A nasal airway inferior turbinate brushing was used to collect airway epithelial cells from these subjects for whole transcriptome sequencing (n=695). Network analyses were performed on all subjects with nasal brushing whole transcriptome sequencing data (n=695) and eQTL analysis was performed on the subset (n=681) with whole genome sequencing generated genotype data.

#### Bulk RNA sequencing of GALA II and ALI Samples

Total RNA was isolated from GALA II subject nasal airway epithelial brushings using the AllPrep DNA/RNA Mini Kit (QIAGEN, Germantown, MD). Whole transcriptome libraries were constructed using the KAPA Stranded mRNA-seq library kit (Roche Sequencing and Life Science, Kapa Biosystems, Wilmington, MA) from 250ng of total input RNA with the Beckman Coulter Biomek FX^P^ automation system (Beckman Coulter, Fullerton, CA) according to the manufacturers protocol. Barcoded libraries were pooled and sequenced using 125bp paired-end reads on the Illumina HiSeq 2500 system (Illumina, San Diego, CA). Bulk RNA-seq data for the nasal and tracheal ALI cultures to measure *ACE2* and *TMPRSS2* levels reported in Figures 3b,c and 4g,h, was generated with KAPA Hyperprep Stranded mRNA-seq library kits (Roche Sequencing and Life Science, Kapa Biosystems, Wilmington, MA) and sequenced with a Novaseq 6000 using 150bp paired end reads.

#### Whole genome sequencing of GALA II Samples

Genomic DNA was extracted from whole blood obtained from GALA II study subjects using the Wizard Genomic DNA Purification kits (Promega, Fitchburg, WI), and DNA was quantified by fluorescent assay. DNA samples were sequenced as part of the Trans□Omics for Precision Medicine (TOPMed) whole genome sequencing (WGS) program^35^. WGS was performed at the New York Genome Center and the Northwest Genomics Center on a HiSeqX system (Illumina, San Diego, CA) using a paired□end read length of 150 base pairs, to a minimum of 30X mean genome coverage. Details on DNA sample handling, quality control, library construction, clustering and sequencing, read processing, and sequence data quality control are described elsewhere^35^. Variant calls were obtained from TOPMed data freeze 8 variant call format files.

#### Experiments using an air-liquid interface, mucociliary culture system

Primary human basal airway epithelial cells were expanded and differentiated at air-liquid interface (ALI) *in vitro* according to established protocols^36^. Paired tracheal ALI cultures were mock-treated or treated with 10 ng/mL IL-13 in media (20 µL apical; 500 µL basolateral) for the final 10 days of differentiation (ALI days 11-21) before harvest and scRNA-seq analysis. In contrast, nasal ALI cultures used for bulk RNA-seq analysis (N = 5 GALA II subjects) were either stimulated with IL-13 for 72h following completion of mucociliary differentiation (25 days) or were infected with human rhinovirus strain A16 for 4 h during the final 24 h of the 28 days of differentiation. Control cultures were only treated with media.

### Preparation of ALI cultures for 10X scRNAseq

Following stimulation experiments involving the tracheal airway epithelial ALI samples, apical culture chambers were washed once with PBS and once with PBS supplemented with dithiothreitol (DTT;10mM), followed by two PBS washes to remove residual DTT. Cold active protease (CAP) solution (*Bacillus licheniformis* protease 2.5 μg/mL, DNase 125 U/mL, and 0.5 mM EDTA in DPBS w/o Ca^2+^Mg^2+^) was added to apical culture chamber and incubated on ice for 10 minutes with mixing every 2.5 minutes. Dissociated cells in CAP solution were added to 500 μL cold FBS, brought up to 5 mL with cold PBS, and centrifuged at 225 x g and 4°C for 5 minutes. The cell pellet was resuspended in 1 mL cold PBS+DTT, centrifuged at 225 x g and 4°C for 5 minutes, and then washed twice with cold PBS. The final cell pellet was resuspended in PBS with 0.04% BSA for single cell gene expression profiling with the 10X Genomics system. Sample capture, cDNA synthesis, and library preparation for 10d IL-13 ALI stimulations was performed using protocols and reagents for 10X Genomics Chromium Single Cell 3’ v3 kit. Single cell libraries were pooled for sequencing on an Illumina NovaSeq 6000.

### Nasal brush 10X scRNA-seq

Nasal brush cells were dissociated from the brush using *Bacillus licheniformis* cold active protease (10mg/ml), EDTA (0.5mM), and EGTA (0.5mM) at 4°C with vortex mixing, followed by enzyme neutralization with FBS. Red blood cell lysis was performed and cells were washed twice in 0.04% BSA/PBS. Cell concentration was adjusted to 400 cells/μL for cell capture of ∼8000 cells using the 10X Genomics Chromium Next GEM Single Cell 3’ reagent kit chemistry. Sample capture, cDNA synthesis, and library preparation was performed following 10X Genomics Chromium Next GEM Single Cell 3’ v3 kit. The single cell library was sequenced on an Illumina NovaSeq 6000.

### QUANTIFICATION AND STATISTICAL ANALYSIS

#### Nasal airway epithelium brushing bulk RNA-seq analysis

##### Preprocessing of RNA-seq data

Raw sequencing reads were trimmed using skewer^37^ (v0.2.2) with the following parameter settings: end-quality=15, mean-quality=25, min=30. Trimmed reads were then aligned to the human reference genome GRCh38 using GSNAP^38^ (v20160501) with the following parameter settings: max-mismatches=0.05, indel-penalty=2, batch=3, expand-offsets=0, use-sarray=0, merge-distant-same-chr. Gene quantification was performed with htseq-count^39^ (v0.9.1) using iGenomes GRCh38 gene transcript model. Variance stabilization transformation (VST) implemented in DESeq2^40^ (v1.22.2) was then carried out on the raw gene count matrix to create a variance stabilized gene expression matrix suitable for downstream analyses.

##### Weighted Gene Co-expression Network Analysis (WGCNA) on GALA II RNA-seq data

To understand what biological mechanisms regulate the variation of nasal airway epithelial gene expression, Weighted Gene Co-expression Network Analysis^41^ (WGCNA) v1.68 was performed on the VST matrix of 17,473 expressed genes. WGCNA analysis is a network-based approach that assumes a scale-free network topology. To adhere to the scale-free assumption of the constructed biological networks, a soft thresholding parameter (ß) value of 9 was chosen based on WGCNA guidelines. Furthermore, minClusterSize was set to 20, deepSplit was set to 2, and pamStage was set to TRUE. A total of 54 co-expression networks were identified and described in Supplementary Table 2. WGCNA networks are referred to by different colors, and two of the these identified networks, saddle brown and tan were found to capture co-expressed genes that underlie T2 inflammation and interferon inflammation, respectively. We hierarchically clustered all subjects based on expression of genes in the saddle brown network and then used the first split in the dendrogram as the basis for assigning individuals to T2-high or T2-low categories (Supplementary Figure 1a). Similarly, we hierarchically clustered subjects using the genes in tan network and then selected the dendrogram branches with the highest tan network expression as interferon-high and the other subjects as interferon-low (Supplementary Figure 2a).

##### Cis-eQTL analysis of nasal RNA-seq data

Cis-expression quantitative trait locus (eQTL) analysis was performed by following the general methodology of the Genotype-Tissue Expression (GTEx) project version 7 protocol^42^, using the nasal RNA-seq data and WGS variant data from 681 GALA II subjects.

Namely, WGS variant data was filtered based on allele frequency (minor allele frequency > 1%) and allele subject count (total number of subjects carrying minor allele ≥ 10). After filtering, 12,590,800 genetic variants were carried forward into the eQTL analysis. For expression data filtering and preparation, we first ran Kallisto^43^ (v0.43.0) to generate transcript per million (TPM) values. We filtered out any genes that did not reach both TPM > 0.1 and raw counts > 6 for at least 20% of our samples. After filtering, 17,039 genes were then TMM normalized using edgeR^44^ (v3.22.3). Finally, we applied an inverse normal transformation into the TMM-normalized expression values to render them suitable for eQTL analysis. To account for global population structure, we ran ADMIXTURE^45^ (v1.3.0) on the genotype data to create five admixture factors. We then ran Probabilistic Estimation of Expression Residuals^46^ (PEER, v1.3) to create 60 PEER factors to utilize as covariates in the eQTL analysis along with admixture estimates, gender, age, body-mass index (BMI), and asthma diagnosis status. To perform cis-eQTL analysis, we utilized a modified version of FastQTL^47^ that was provided by the GTEx project. Furthermore, we performed stepwise regression analysis to identify independent eQTL variants using QTLTools^48^ (v1.1). Allelic Fold Change (A_FC_) of the eQTL variant is computed using the aFC python script^49^.

##### Virus identification and quantification from bulk RNA-Seq data

To identify individuals with asymptomatic virus infection at the time of sample collection, viral genomic sequences were recovered from bulk RNA-seq data using a modified version of the Virus Finder 2.0 (VF2) pipeline^50^. A custom respiratory virus reference database comprising >600k sequences was employed to improve specificity. Using VF2, viral reads were garnered by removing human reads using Bowtie2^51^ (default settings) and selecting viral reads using BLAT^52^ (minIdentity=80); contigs were assembled using Trinity^53^; short (<200 bp) or low complexity (DUST score < 0.07) contigs and contigs matching the human genome at a BLAST^54^ e-value <0.05 were discarded; the remaining contigs were classified using BLAST (e-value <0.05); read counts were obtained by read mapping using BLAT (minIdentity=98). Of the 468 distinct viral reference sequences detected by VF2, 7 were identified as erroneous and removed. The remaining 461 matches were manually assigned viral serotypes and the results aggregated with R.

##### Defining CoV and HRV infected groups and associated analysis

To ensure we selected subjects that were experiencing an active host response to a CoV infection, we examined the distribution of viral reads for the 18 CoV^+^ infected subjects. We observed a clear break between the 7 subjects with the lowest viral read counts (<3,000 reads) and 11 subjects with the highest viral read counts (>60,000 reads). Therefore, we selected these 11 highly infected subjects for analysis of host responses to CoV infection. To generate a similar infection-control group, composed of subjects highly infected with a different virus species, we examined the 67 HRV infected subjects in GALA, enforcing a comparable lower bound of viral reads as with CoV, adjusting for the smaller HRV genome size. Specifically, HRV genomes are ∼7,000 base pairs, whereas CoV genomes are ∼30,000 base pairs, making the HRV genome ∼25% of the size of the CoV genome. Therefore, we selected a cutoff of 15,000 viral reads for subjects to be included in the HRV^+^ highly infected group. Therefore, we selected a cutoff of 15,000 viral reads for subjects to be included in the HRV^+^ highly infected group (n=37) analyzed in Figure 6. All non-infected subjects (n=571), based on the Virus Finder analysis described above, were used as comparison group for the CoV^+^ and HRV^+^ groups.

In performing the CoV^+^ and HRV^+^ transcriptome-wide differential expression analyses, to account for the class imbalance of this experiment, log_2_ count-normalized expression values in units of counts per million (calculated using edgeR v3.28.0) were passed to the function arrayWeights function in the limma^55^ R package (3.42.0). limma-voom was then used to perform differential expression analysis on the count normalized expression values between the CoV^+^ and uninfected groups, as well as between the HRV^+^ and uninfected groups, controlling for age, gender, and asthma diagnosis status. Genes were required to have an FDR adjusted p-value < 0.05, and an absolute log_2_FC > 0.5 to be considered significant. Based on these cutoffs, genes were classified as being shared if they were significant in both comparisons, or as CoV^+^–specfic or HRV^+^– specific if significant in only one comparison.

##### Gene set enrichment analysis

To investigate enriched pathways within WGCNA networks (see Figure 2a) or within genes differentially expressed in CoV^+^ and/or HRV^+^ infected subject groups (see Figure 6c and 6c), we used Enrichr^56^ to test for gene overrepresentation of network genes within a panel of annotated gene databases (Gene Ontology [GO] Biological Process [BP] 2018, GO Molecular Function [MF] 2018, GO Cellular Component [CC] 2018, Kyoto Encyclopedia of Genes and Genomes [KEGG] 2019 Human, and Reactome 2016). For cell type enrichments within WGCNA networks reported in Figure 2a, we tested for overrepresentation of network genes within gene marker sets (FDR < 0.05) for each of 35 epithelial and immune cell types inferred using scRNA-seq of human lung tissue^57^.

For the plots in Figure 6f-g, transcriptomic data for 11 flow sorted immune cell populations were obtained from GEO experiments GSE3982 and GSE22886 and then batch corrected using the ComBat^58^ function from the SVA R package (v3.34.0). limma was then used to perform differential expression analysis between each cell type and all the rest in order to obtain gene log_2_FC values for each cell type with which to rank order the genes. Gene set enrichment analysis (GSEA) was then used to test for association between upregulated genes in the shared, CoV^+^–specific, and HRV^+^–specific gene sets and each of the cell types, based on the cell type-specific ordered gene lists. GSEA was carried out using the FGSEA R package (v1.12.0).

##### Canonical pathway analysis

We used QIAGEN’s Ingenuity Pathway Analysis (IPA) program (v01-16; content version: 51963813, release 2020-03-11) to investigate canonical pathways and upstream regulators that were significantly enriched in one or both of the upregulated CoV^+^-specific or HRV^+^-specific gene sets.

#### Analysis of scRNA-seq data from the nasal epithelial brushing

Initial processing of 10X scRNA-seq data, including cell demultiplexing, alignment to the human genome GRCh38, and UMI-based quantification was performed with Cell Ranger (version 3.0). Since the nasal brushing sample contains both epithelial and immune cell populations that have distinct expression profiles (e.g.: Immune cell types express far fewer genes compared to epithelial cell types), clustering and cell type identification were done in two stages: 1) an initial clustering with a less stringent filter to identify major epithelial and immune cell clusters was performed, 2) cells were reclustered with different independent filtering criteria for epithelial and immune cell types. All these analyses were performed using Seurat^59^ R package (v3.0).

In the first stage, we removed cells with fewer than 100 genes detected or cells with greater than 25% mitochondrial reads. Additionally, to remove possible doublets, we removed cells with higher than 6,000 genes detected and/or more than 20,000 UMIs. Lowly expressed genes (detected in fewer than 4 cells) were also removed. We then performed normalization using SCTransform^60^ and ran PCA on the top 5000 highly variable normalized genes. Clustering analysis was performed on the top 20 PCs using a shared nearest neighbor (SNN) based SLM^61^ algorithm with the following parameter settings: resolution=0.8, algorithm=3. The single cell expression profiles were visualized via embedding into two dimensions with UMAP^62^ (Uniform Manifold Approximation and Projection), resulting in the identification of 11,157 epithelial cells and 229 immune cells based on known cell type signatures.

In the second stage, we retained all the immune cells but removed epithelial cells with fewer than 1,000 detected genes. After this filtering, a combined 8,291 epithelial and immune cells were then normalized as in the first stage. Clustering analysis performed on the top 30 PCs with parameters (resolution=0.4, algorithm=1, k.param=10) identified 15 clusters. We then ran differential expression analysis using a Wilcoxon test implemented in Seurat’s “FindMarkers” function to help with cell type identification. Based on these cluster marker lists, two clusters were merged into a single secretory cluster, another two clusters were merged into a single ciliated cluster, and a final two clusters were combined as “indeterminate,” based on the lack of defining marker genes for these clusters. Through this merging process, we arrived at 8 epithelial and 3 immune cell populations (Figure 1a, Supplementary Table 1)

#### Analysis of bulk RNA-seq data from IL-13 and HRV infected ALI nasal airway epithelial cultures

Raw sequencing reads were trimmed using skewer with the following parameter settings: end-quality=15, mean-quality=25, min=30. Trimmed reads were then aligned to the human reference genome GRCh38 using HISAT2^63^ (v2.1.0) using default parameter settings. Gene quantification was performed with htseq-count using the GRCh38 Ensembl v84 gene transcript model. After removing mitochondrial, ribosomal, and lowly expressed genes (those not expressed in at least two samples), we carried out differential expression analyses between paired IL-13-stimulated and control samples (N = 5 donors) and between paired HRV-infected and control samples (N = 5 donors) using the DESeq2 R package (v1.22.2).

#### Analysis of scRNA-seq data from 10 day IL-13-stimulated and control tracheal cell ALI cultures

As with the nasal brushing scRNA-seq data, 10X scRNA-seq data from ALI cultures grown from a single tracheal donor that were either mock- or IL-13 stimulated for 10 days were pre-processed using Cell Ranger (version 3.0, 10X Genomics). To safeguard against doublets, we removed all cells with gene or UMI counts exceeding the 99th percentile. We also removed cells expressing fewer than 1,500 genes or for which > 30% of genes were mitochondrial (genes beginning with *MTAT, MT*-, *MTCO, MTCY, MTERF, MTND, MTRF, MTRN, MRPL*, or *MRPS*), resulting in a total of 6,969 cells (2,715 IL-13-stimulated and 4,254 controls). After removing mitochondrial, ribosomal (*RPL* and *RPS*), and very lowly expressed genes (expressed in < 0.1% of cells), we integrated expression data from IL-13 and control cells using the dataset integration approach in Seurat^64^. For the integration analysis, we used the top 30 dimensions from a canonical correlation analysis (CCA) based on SCTransform normalized expression of the top 3,000 most informative genes across the two datasets, where “informativeness” was defined by gene dispersion (i.e., the log of the ratio of expression variance to its mean) across cells, calculated after accounting for its relationship with mean expression. We then carried out principle component analysis (PCA) on the integrated dataset and used the top 20 components for clustering and visualization. We used SNN (Louvain algorithm, resolution=0.23, k.param=10) to cluster the integrated cells into 11 populations, which we visualized in two dimensions using UMAP (see Figure 3d). These clusters were assigned cell type labels based their most upregulated genes, which were identified by carrying out differential expression analysis between each cluster and all others using Seurat’s logistic regression (LR) test, in which cell treatment was included as a latent variable.

## DATA AVAILABILITY

All raw and processed RNA-seq data used in this study are currently being deposited in the National Center for Biotechnology Information/Gene Expression Omnibus (GEO).

## CODE AVAILABILITY

## Acknowledgements

This work was supported by NIH grants (MAS) U01 HL138626, R01 HL135156, R01 MD010443, R01 HL128439, P01 HL132821, P01 HL107202, R01 HL117004, and DOD Grant W81WH-16-2-0018.

The Genes-Environments and Admixture in Latino Americans (GALA II) Study and E.G.B. were supported by the Sandler Family Foundation, the American Asthma Foundation, the RWJF Amos Medical Faculty Development Program, the Harry Wm. and Diana V. Hind Distinguished Professor in Pharmaceutical Sciences II, the National Heart, Lung, and Blood Institute (NHLBI) [R01HL117004, R01HL128439, R01HL135156, X01HL134589]; the National Institute of Environmental Health Sciences [R01ES015794]; the National Institute on Minority Health and Health Disparities (NIMHD) [P60MD006902, R01MD010443], the National Human Genome Research Institute [U01HG009080] and the Tobacco-Related Disease Research Program [24RT-0025, 27IR-0030]. MJW was supported by the NHLBI [K01HL140218]. Burchard NIH Support: T32 GM007546, U01 HL138626, R01 128439, R01 HL141992, R01 HL141845.

Whole genome sequencing (WGS) for the Trans-Omics in Precision Medicine (TOPMed) program was supported by the National Heart, Lung and Blood Institute (NHLBI). WGS for “NHLBI TOPMed: Gene-Environment, Admixture and Latino Asthmatics Study” (phs000920) was performed at the New York Genome Center (3R01HL117004-02S3) and the University of Washington Northwest Genomics Center (HHSN268201600032I). Centralized read mapping and genotype calling, along with variant quality metrics and filtering were provided by the TOPMed Informatics Research Center (3R01HL-117626-02S1; contract HHSN268201800002I). Phenotype harmonization, data management, sample-identity QC, and general study coordination were provided by the TOPMed Data Coordinating Center (3R01HL-120393-02S1, U01HL-120393, contract HHSN268201800001I). We gratefully acknowledge the studies and participants who provided biological samples and data for TOPMed.

WGS of part of GALA II was performed by New York Genome Center under The Centers for Common Disease Genomics of the Genome Sequencing Program (GSP) Grant (UM1 HG008901). The GSP Coordinating Center (U24 HG008956) contributed to cross-program scientific initiatives and provided logistical and general study coordination. GSP is funded by the National Human Genome Research Institute, the National Heart, Lung, and Blood Institute, and the National Eye Institute.

The authors wish to acknowledge the following GALA II study collaborators: Shannon Thyne, UCSF; Harold J. Farber, Texas Children’s Hospital; Denise Serebrisky, Jacobi Medical Center; Rajesh Kumar, Lurie Children’s Hospital of Chicago; Emerita Brigino-Buenaventura, Kaiser Permanente; Michael A. LeNoir, Bay Area Pediatrics; Kelley Meade, UCSF Benioff Children’s Hospital, Oakland; William Rodríguez-Cintrón, VA Hospital, Puerto Rico; Pedro C. Ávila, Northwestern University; Jose R. Rodríguez-Santana, Centro de Neumología Pediátrica; Luisa N. Borrell, City University of New York; Adam Davis, UCSF Benioff Children’s Hospital, Oakland; Saunak Sen, University of Tennessee.

The authors acknowledge the families and patients for their participation and thank the numerous health care providers and community clinics for their support and participation in GALA II. In particular, the authors thank the recruiters who obtained the data: Duanny Alva, MD; Gaby Ayala-Rodríguez; Lisa Caine, RT; Elizabeth Castellanos; Jaime Colón; Denise DeJesus; Blanca López; Brenda López, MD; Louis Martos; Vivian Medina; Juana Olivo; Mario Peralta; Esther Pomares, MD; Jihan Quraishi; Johanna Rodríguez; Shahdad Saeedi; Dean Soto; and Ana Taveras.

The content is solely the responsibility of the authors and does not necessarily represent the official views of the National Institutes of Health.

## Notes

### Competing Interest Statement

The authors have declared no competing interest.

