## Supplementary Figure legend for "Type 2 and interferon inflammation strongly regulate SARS-CoV-2 related gene expression in the airway epithelium"

*Authors/Affiliations*

Satria P. Sajuthi^1#^, Peter DeFord^1#^, Nathan D. Jackson^1^, Michael T. Montgomery^1^, Jamie L. Everman^1^, Cydney L. Rios^1^, Elmar Pruesse^1^, James D. Nolin^1^, Elizabeth G. Plender^1^, Michael E. Wechsler^2^, Angel CY Mak^3^, Celeste Eng^3^, Sandra Salazar^3^, Vivian Medina^4^, Eric M. Wohlford^3,5^, Scott Huntsman^3^, Deborah A. Nickerson^6,7,8^, Soren Germer^9^, Michael C. Zody^9^, Gonçalo Abecasis^10^, Hyun Min Kang^10^, Kenneth M. Rice^11^, Rajesh Kumar^12^, Sam Oh^3^, Jose Rodriguez-Santana^4^, Esteban G. Burchard^3,13^, Max A. Seibold^1,14,15^*

^1^Center for Genes, Environment, and Health, National Jewish Health, Denver, CO, 80206 USA; ^2^Department of Medicine, ^14^Department of Pediatrics, National Jewish Health, Denver, CO, 80206 USA; ^15^Division of Pulmonary Sciences and Critical Care Medicine, University of Colorado-AMC, Aurora, CO, 80045 USA, ^3^Department of Medicine, ^5^Division of Pediatric Allergy and Immunology, ^13^Department of Bioengineering and Therapeutic Sciences University of California San Francisco, San Francisco, CA; ^6^Department of Genome Sciences, University of Washington, Seattle, WA, USA, ^7^Northwest Genomics Center, Seattle, WA, USA ^8^Brotman Baty Institute, Seattle, WA, USA; ^9^New York Genome Center, NYC, New York; ^10^Center for Statistical Genetics, University of Michigan, Ann Arbor, MI, USA; ^11^Department of Biostatistics, University of Washington, Seattle, WA, USA; ^4^Centro de Neumología Pediátrica, San Juan, Puerto Rico; ^12^Ann and Robert H. Lurie Children's Hospital of Chicago, Department of Pediatrics, Northwestern University, Chicago, Ill ^#^these authors contributed equally to this work, *correspondence:

*Supplement Figure Legends*

**Figure S1. Identification of T2-high subjects and association with clinical traits**

(a) Clustering of subjects by expression of T2 biomarker network (saddle brown) genes identifies T2-high and T2-low groups. Heatmap of the T2 biomarker network genes across all clustered subjects. The colorbars at the top of the heatmap correspond to the T2 clusters. The blue and red bars identify T2-low and T2-high subjects, respectively.

(b) Distribution of T2-status assignments based on asthma status. T2 status is associated with asthma status.

(c-e) Box plots of atopy associated clinical variable levels by T2-status. P-values are generated from a Wilcoxon-rank-sum test.

(c) Fractional exhaled nitric oxide in parts per billion (FeNO) by T2-status.

(d) Total serum IgE levels by T2-status.

(e) Whole blood eosinophil levels by T2-status.

**Figure S2. Identification of interferon-high subjects and association with viral infection status**

(a) Clustering of subjects by expression of interferon response network genes (tan) identifies interferon-high and interferon-low groups. Heatmap of the tan network genes across all clustered subjects. The colorbars at the top of the heatmap correspond to the interferon clusters. The blue and red bars identify interferon-low and interferon-high subjects, respectively.

(b) Distribution of interferon-status assignments based on viral infection status. Interferon-high status is associated with viral infection status.

**Figure S3. Additional independent eQTLs for *TMPRSS2***

(a) Boxplot of *TMPRSS2* expression by genotype for eQTL variant rs74659079.

(b) Boxplot of *TMPRSS2* expression by genotype for eQTL variant rs2838057.

**Figure S4. UMAP of scRNA-seq data from chronic IL-13 culture experiment with cells colored by IL-13 or mock treatment status**

Blue and red cells come from cultures stimulated with either mock or IL13 stimulus, respectively.

**Figure S5. Scatterplot of Log_2_FC values for the union of DEGs for the CoV and HRV infection response analyses**

Scatterplot of the log fold changes in expression for virus infected groups (CoV or HRV) vs. the uninfected group shows high correlation in fold changes of DEGs for each virus. Points represent DEGs and are colored by whether they were significant for CoV+ only (yellow), HRV+ only (red), or both analyses (blue).
