## Supplementary figures and images for "Type 2 and interferon inflammation strongly regulate SARS-CoV-2 related gene expression in the airway epithelium"

### Supplementary Figure 1

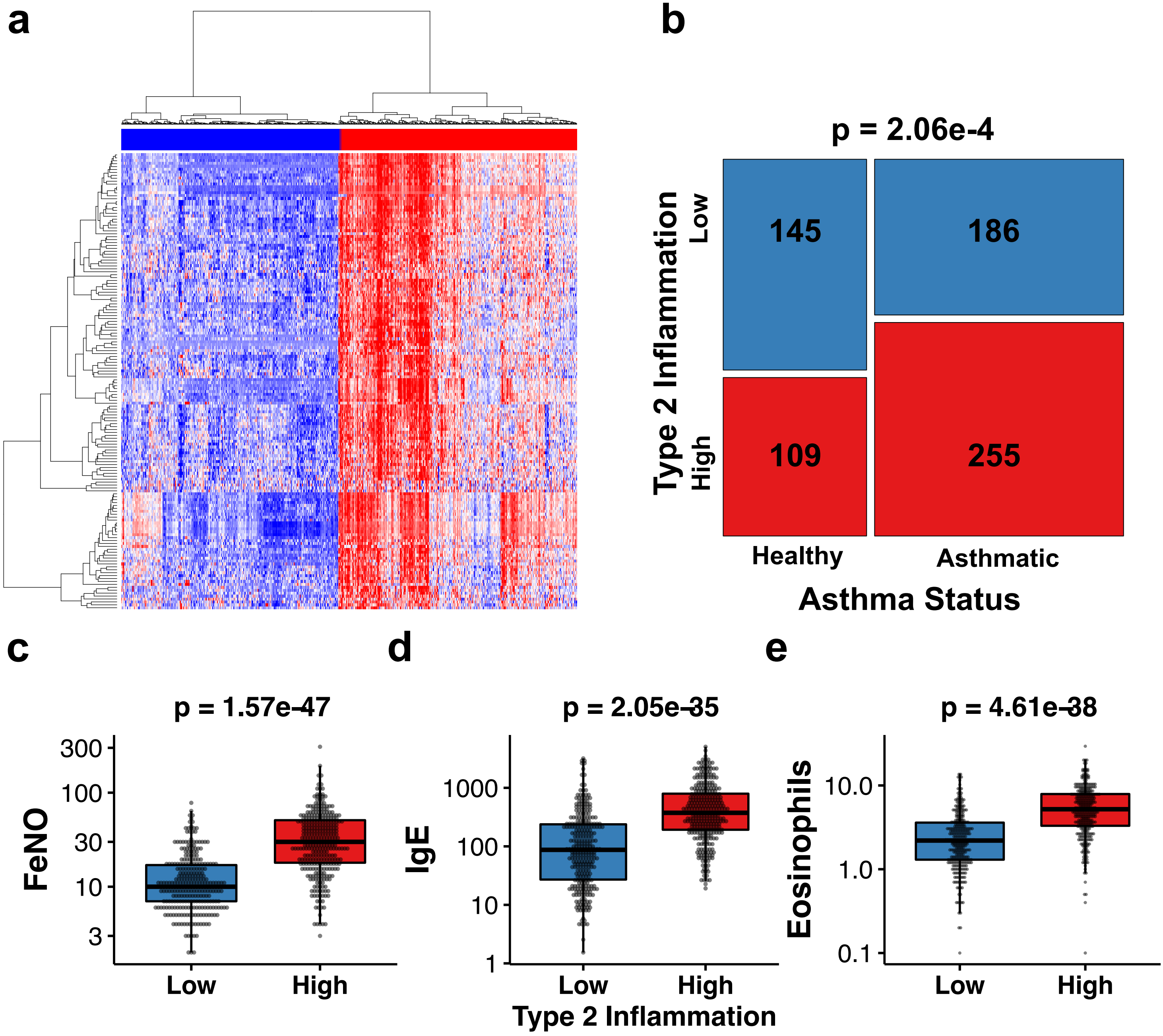

### Supplementary Figure 2

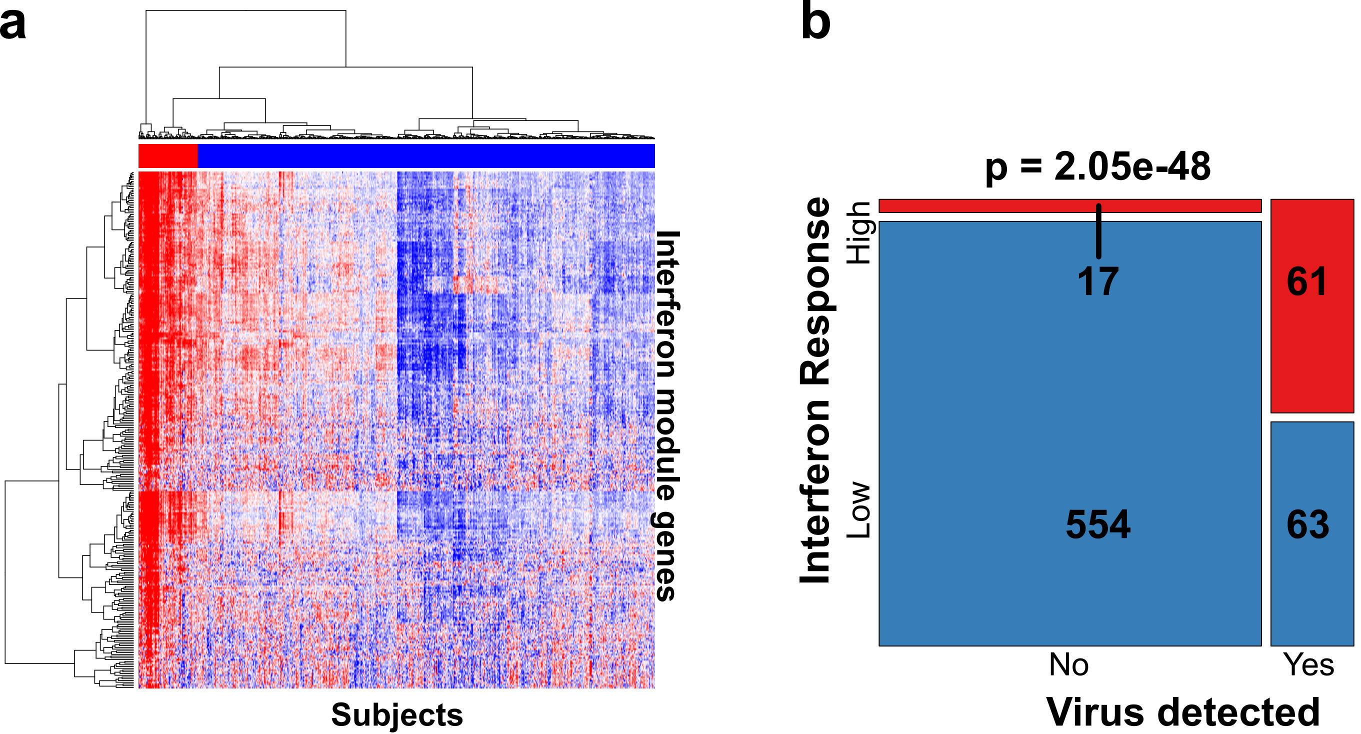

### Supplementary Figure 3

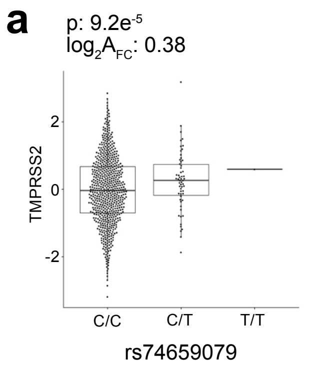

### Supplementary Figure 4

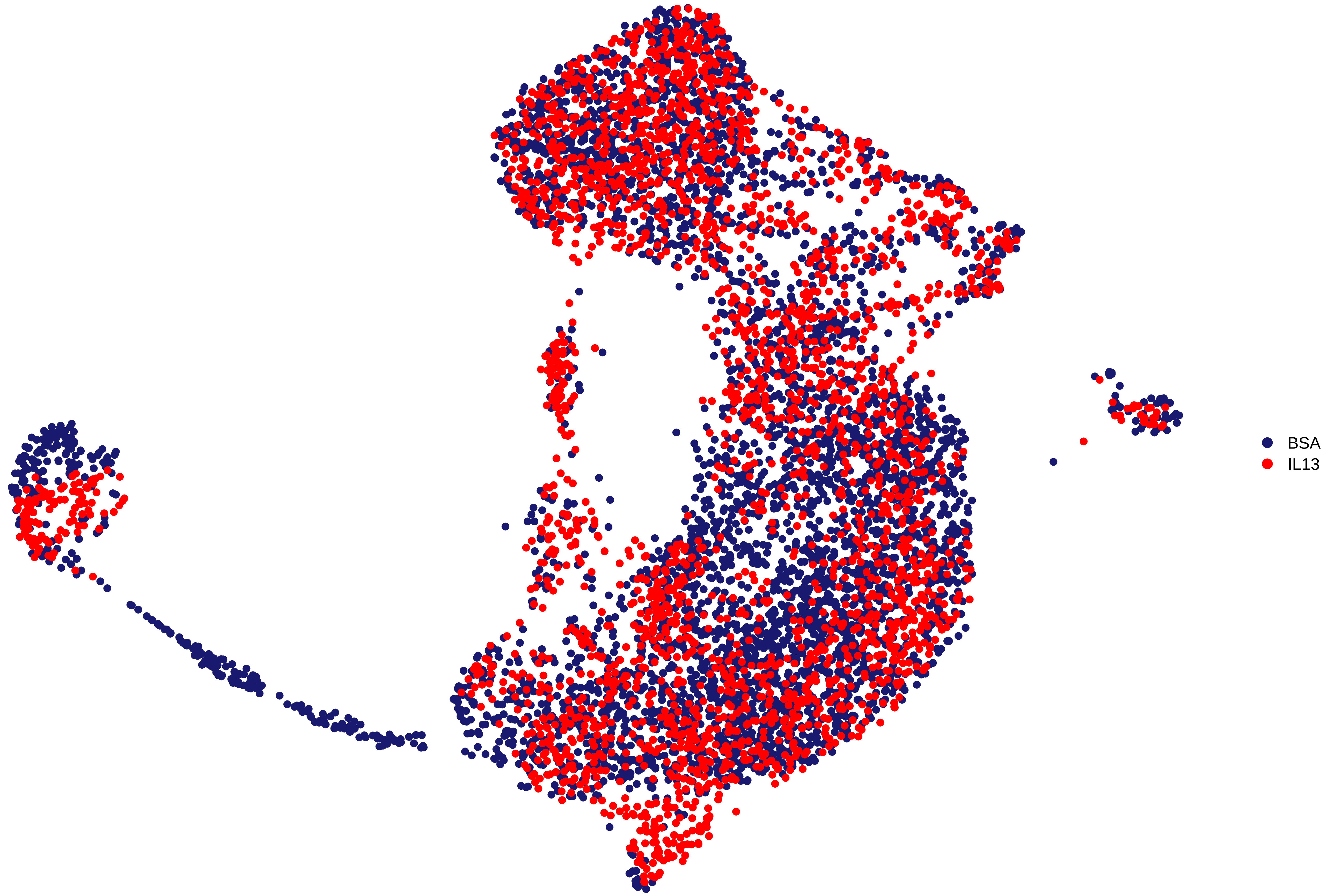

### Supplementary Figure 5

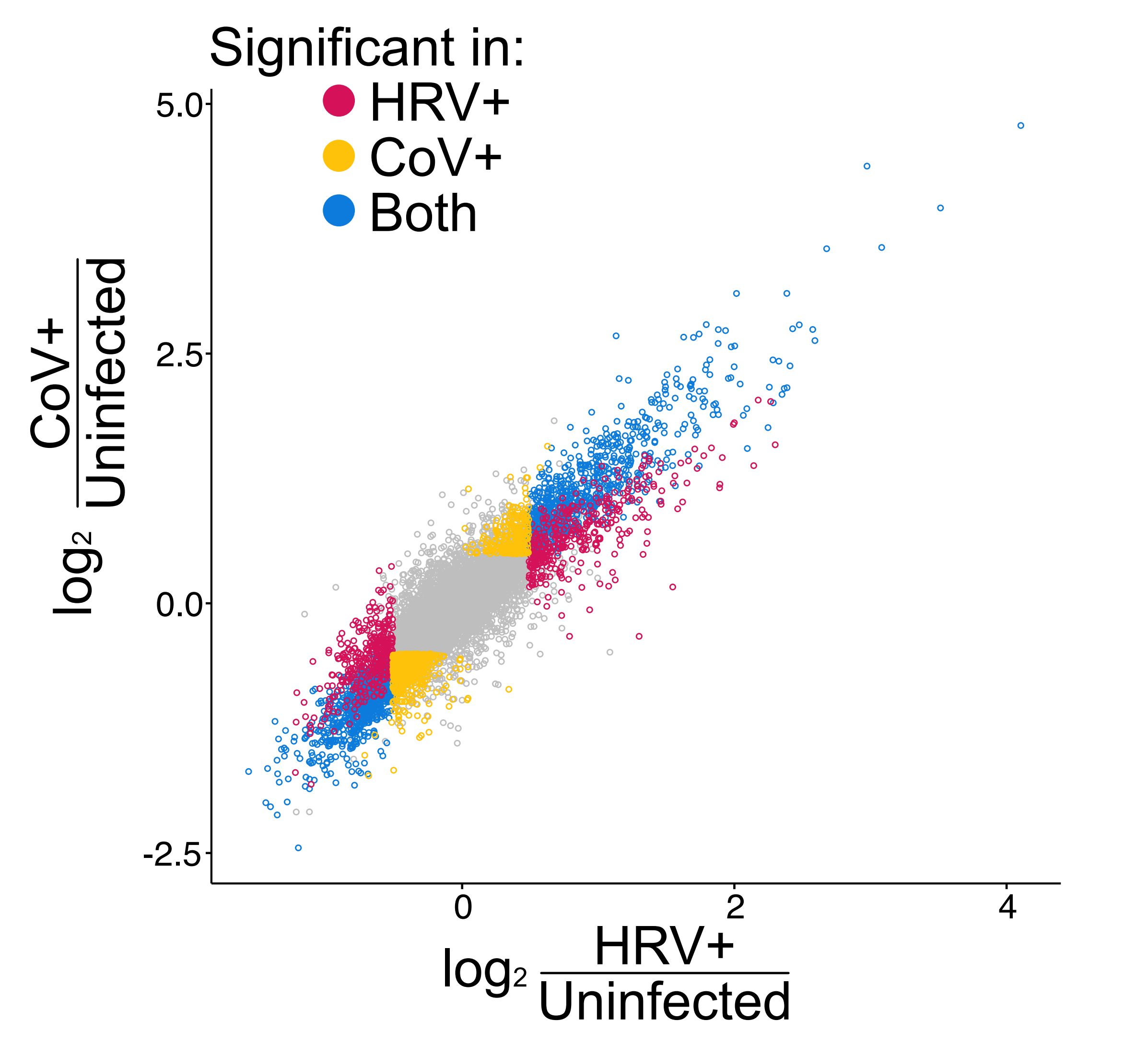
